# Liana optical traits increase tropical forest albedo and reduce ecosystem productivity

**DOI:** 10.1101/2021.06.08.447067

**Authors:** Félicien Meunier, Marco D. Visser, Alexey Shiklomanov, Michael C. Dietze, J. Antonio Guzmán Q., Arturo Sanchez-Azofeifa, Hannes P. T. De Deurwaerder, Sruthi M. Krishna Moorthy, Stefan A. Schnitzer, David C. Marvin, Marcos Longo, Liu Chang, Eben N. Broadbent, Angelica M. Almeyda Zambrano, Helene Muller-Landau, Matteo Detto, Hans Verbeeck

## Abstract

Lianas are a key growth form in tropical forests. Their lack of self-supporting tissues and their vertical position on top of the canopy make them strong competitors of resources. A few pioneer studies have shown that liana optical traits differ on average from those of colocated tree. Those trait discrepancies were hypothesized to be responsible for the competitive advantage of lianas over trees. Yet, in the absence of reliable modelling tools, it is impossible to unravel their impact on the forest energy balance, light competition and on the liana success in Neotropical forests. To bridge this gap, we performed a meta-analysis of the literature to gather all published liana leaf optical spectra, as well as all canopy spectra measured over different levels of liana infestation. We then used a Bayesian data assimilation framework applied to two radiative transfer models (RTMs) covering the leaf and canopy scales to derive tropical tree and liana trait distributions, which finally informed a full dynamic vegetation model. According to the RTMs inversion, lianas grew thinner, more horizontal leaves with lower pigment concentrations. Those traits made the lianas particularly efficient at light interception and completely modified the forest energy balance and its carbon cycle. While forest albedo increased by 14% in the shortwave, light availability was dramatically reduced in the understory (−30% of the PAR radiation) and soil temperature decreased by 0.5°C. Those liana-specific traits were also responsible for a significant reduction of tree (−19%) and ecosystem (−7%) gross primary productivity (GPP) while lianas benefited from them (their GPP increased by +27%). This study provides a novel mechanistic explanation to the increase in liana abundance, new evidence of the impact of structural parasitism on forest functioning, and paves the way for the evaluation of the large-scale impacts of woody vines on forest biogeochemical cycles.

## Introduction

Terrestrial ecosystems are a key component of the Earth’s carbon cycle as they are responsible for a yearly uptake of about 60 GtC (Beer et al. 2010) and store about 860 GtC worldwide (Pan et al. 2011), about 50% of which is located in the tropics (Brinck et al. 2017; Avitabile et al. 2016). Forests in general, and tropical ecosystems in particular, also profoundly regulate the global energy budget by mediating the exchange of energy and moisture between the land and the atmosphere (Fischlin et al. 2007; Piao et al. 2020; Spracklen et al. 2018). Through succession, land-use, management, and vegetation dynamics, the ratio of back-reflected solar radiation to the total received (*i.e*. albedo) may radically change (Bonan 2008).

Forest albedo is controlled by the optical properties and geometric arrangement of leaf and wood tissues, and, in open canopies, also by the contribution of soil reflectivity. All together, these forest features determine what fraction of incident light penetrates in the canopy, to what depth, where light is absorbed, and how much is reflected back to the atmosphere (Asner 2008). Light interception and distribution within the canopy not only regulates photosynthesis but also influences long term processes like recruitment, competition among and within species, and phenology (Bonan 2019). In other words, radiative transfer is a key-process for all plants in the ecosystem (Yuan et al. 2016) and its accurate representation in vegetation models is critical to represent ecosystem functioning (Fisher et al. 2018).

While radiative transfer in canopies has been studied for decades (Jacquemoud et al. 2000), as of today there has been little focus on its contribution to uncertainty in dynamic global vegetation models (Viskari et al. 2019). However, as light is often the limiting resource in dense tropical canopies, it plays a critical role for plant growth and development, often driving intra- and interspecies competition and succession (Bongers and Sterck 1998; Poorter et al. 2003). Moreover, in the context of global warming, a change of tropical ecosystem albedo might have important feedbacks on the regional climate (Piao et al. 2020).

Lianas are woody vines that are abundant in tropical ecosystems (Schnitzer 2005) where they act as structural parasites of the forest (Stevens 1987). They climb up the stems of other plants to reach the top of the canopy from which they compete for light and progressively displace a significant fraction of tree leaf biomass with their own (Schnitzer et al. 2005; Selaya and Anten 2008; Kazda and Salzer 2000). In tropical forests, lianas play a key role in vegetation dynamics as they represent on average 15% of the woody stems (Schnitzer and Bongers 2002; Dewalt et al. 2014) and contribute to 9-31% of the total leaf area (van der Heijden et al. 2013).

Previous studies have demonstrated that the leaf spectral signature can significantly differ between tropical trees and lianas (Castro-Esau et al. 2004; Kalacska et al. 2007; Sánchez-Azofeifa et al. 2009; Guzmán et al. 2018). These findings are consistent with multiple observations of growth form level differences in leaf biochemical traits such as chlorophyll, carotenoid, and water content (Asner and Martin 2012) and a few seminal studies suggesting an increase of albedo in canopies with high liana coverages (Sánchez-Azofeifa and Castro-Esau 2006; Marvin et al. 2016; Kalacska et al. 2007). Liana leaves, because of their position in the canopy (often at the top) and their contrasting biochemical properties as compared to trees, could profoundly impact forest functioning by modifying the forest energy cycle. We argue that a significant part of the changes of albedo observed by remote sensing in tropical forests (Doughty et al. 2018; Piao et al. 2020) might be due to liana infestation variability. This hypothesis is especially relevant to test in the context of increasing liana abundance observed in the Neotropics (Phillips 2002; Schnitzer and Bongers 2011), which could aggravate the impact of lianas on radiative transfer of tropical forests in the near future.

Despite their potential impact on forest biogeochemical cycles, lianas have generally been ignored in vegetation models and remote sensing products (Krishna Moorthy 2019; Verbeeck and Kearsley 2016). However, the recent implementation of the lianescent growth form in the Ecosystem Demography model (di Porcia e Brugnera et al. 2019; Meunier et al. 2020) now supports evaluating the role of lianas on the energy budget of tropical ecosystems under changing environmental conditions. Because vegetation models combine the biophysics of land surface models with vegetation demography and biogeochemistry (Purves and Pacala 2008; Fisher et al. 2014; Dietze et al. 2014), they are excellent candidates for studying and projecting the impacts of climate change and resulting forest composition evolution (such as liana proliferation) on the radiation profile and forest albedo (Fisher et al. 2018).

The objective of this study is to investigate how lianas influence the energy budget and the biogeochemical cycles of tropical forests by altering light interception. We hypothesized that liana-specific optical traits (i) result in an increased albedo and a decreased understorey light availability in heavily infested forest patches and (ii) are responsible for earlier observations of liana-induced ecosystem and tree declined productivity. If these hypotheses are confirmed, the impact of liana proliferation might be more complex than previously expected as earlier studies have limited their scope to the contribution of lianas to the carbon cycle.

## Material and Methods

To estimate the impact of liana-specific traits on the radiative transfers of tropical forests, we first assembled a database of liana leaf-level reflectance measurements from literature, along with stand-level reflectance spectra covering different levels of liana infestation. We assimilated the collected spectra to derive leaf biochemical and canopy structural properties using leaf-level (PROSPECT-5) and stand-level (ED-RTM) radiative transfer models and estimate how those traits diverge between tropical trees and lianas. We then ran simulations of a process-based vegetation model (ED2.2), in which lianas were in turn characterized by liana or tree parameter distributions. We validated the principal findings of this study with independent experimental datasets (WorldView-3 and GatorEye UAV-LiDAR). The overall workflow of this study is illustrated in Figure 1.

**Figure 1:**
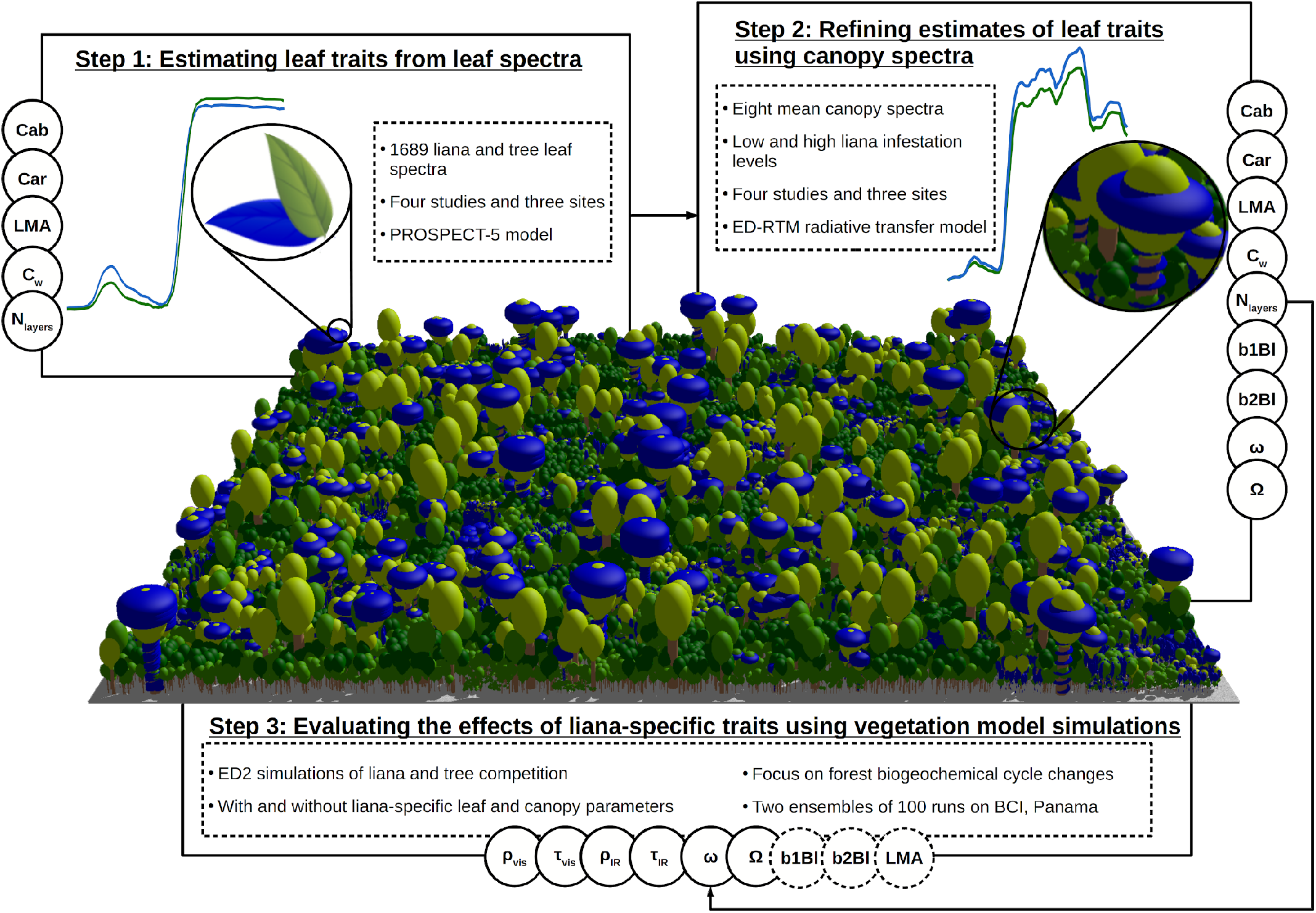
Workflow of the study as divided in three steps. Observed leaf spectra of both lianas and trees were assimilated to optimize leaf biochemical parameters of those PFTs through PROSPECT-5 simulations (Step 1). Leaf optical and biochemical traits were further calibrated in a radiative transfer model (ED-RTM) together with liana and tree canopy parameters through the assimilation of canopy reflectance spectral data under low and high liana infestation levels (Step 2). The resulting parameter posterior distributions then served to evaluate the impact of liana leaf parameters in a vegetation model (ED2.2) in simulations with and without liana-specific leaf and canopy parameters. In these runs, liana LAI-related parameters (b1Bl, b2Bl, and LMA) were systematically sampled from liana parameter distributions to conserve a similar ecosystem LAI. In steps 1 and 2, all parameters indicated on the side were optimized to fit observational data within a Bayesian framework. Throughout the manuscript, lianas (and liana-rich forest stands) are consistently represented in blue.

## Model description

### The PROSPECT-5 model

The PROSPECT-5 model simulates the spectral reflectance and transmittance of a leaf over a large wavelength range covering the visible (400-700 nm), near infrared (700-1400 nm), and shortwave infrared (1500-2500 nm) at the nanometer resolution. PROSPECT-5 represents leaves as stacks of partially reflective/transparent layers and is based on five biochemical and structural properties, namely the total chlorophyll (*Cab*) and Carotenoid (*Car*) contents, the number of stacked elementary homogeneous layers (*N*_*layers*_), the leaf equivalent water thickness (*C*_*w*_) and its dry matter content (*LMA*) (Feret et al. 2008). Wavelength-specific reflectivity and transmissivity coefficients are then calculated as a weighted linear combination of the empirically-calibrated absorption spectra for leaf pigments, water, and dry matter (Feret et al. 2008). Table 1 summarises the parameters discussed in this study, together with their units and their description.

**Table 1:** Leaf biochemical and canopy structural traits of the radiative transfer models used in this study, together with their prior distributions for lianas and trees as well as the posterior medians. The prior parameters column provides the minimum (a) and maximum (b) values for the uniform distributions, or the mean (a) and the standard deviation (b) for the normal distributions.

| Abbreviation | Units | Description | PFT | Prior | Prior parameters | Posterior median [95% CI]<sup>†</sup> |
| --- | --- | --- | --- | --- | --- | --- |
| Cab | $\mu\text{g cm}^{-2}$ | Leaf chlorophyll area density | Liana | Uniform | a = 0, b = 150 | 45.5 [45.3 45.6] |
|  |  |  | Tree | Uniform | a = 0, b = 150 | 56.6 [56.4 56.8] |
| Car | $\mu\text{g cm}^{-2}$ | Leaf carotenoid area density | Liana | Uniform | a = 0, b = 50 | 15.9 [15.8 6.0] |
|  |  |  | Tree | Uniform | a = 0, b = 50 | 20.6 [20.5 20.8] |
| $C_w$ | cm | Equivalent water thickness | Liana | Uniform | a = 0, b = 0.1 | 0.0152 [0.0151 0.152] |
|  |  |  | Tree | Uniform | a = 0, b = 0.1 | 0.0199 [0.0197 0.0200] |
| LMA<sup>*</sup> | $\text{kg m}^{-2}$ | Leaf dry matter content per unit area | Liana | Uniform | a = 0.01, b = 1 | 0.064 [0.064 0.064] |
|  |  |  | Tree | Uniform | a = 0.01, b = 1 | 0.087 [0.086 0.088] |
| $N_{\text{layers}}$ | - | Effective number of mesophyll layers | Liana | Uniform | a = 1.01, b = 5 | 1.76 [1.76 1.77] |
|  |  |  | Tree | Uniform | a = 1.01, b = 5 | 2.06 [2.05 2.06] |
| b1Bl | $\text{kg plant}^{-1} \text{cm}^{-b2Bl}$ | Leaf biomass allometry intercept | Liana | Normal | a = 0.049, b = 0.010 | 0.049 [0.049 0.049] |
|  |  |  | Tree | Normal | a = 0.020, b = 0.005 | 0.020 [0.020 0.020] |
| b2Bl | - | Leaf biomass allometry slope | Liana | Normal | a = 1.89, b = 0.2 | 1.89 [1.89 1.90] |
|  |  |  | Tree | Normal | a = 1.85, b = 0.2 | 1.85 [1.85 1.85] |
| $\omega$ | - | Leaf orientation factor | Liana | Uniform | a = -0.5, b = 0.5 | 0.33 [0.32 0.34] |
|  |  |  | Tree | Uniform | a = -0.5, b = 0.5 | -0.42 [-0.42 -0.41] |
| $\Omega$ | - | Leaf clumping factor | Liana | Uniform | a = 0.4, b = 1 | 0.72 [0.71 0.72] |
|  |  |  | Tree | Uniform | a = 0.4, b = 1 | 0.48 [0.48 0.48] |

<sup>\*</sup>Equivalently referred to as $C_m$ (the units of which are $\text{g cm}^{-2}$ in PROSPECT-5).
<sup>†</sup>Posteriors from all studies/sites.

### The Ecosystem Demography model, version 2

#### Overview of the vegetation model

The Ecosystem Demography model version 2.2 (ED2.2) is a vegetation model that simulates forest biophysical and physiological cycles, and accounts for horizontal and vertical heterogeneities of the ecosystem (Longo et al. 2019a). Carbon, water, and energy cycles are solved for individual forest patches, each containing zero, one, or several plant cohorts. Patches are defined as areas of the forest with similar age (*i.e*. disturbance history) while cohorts are groups of plants in a patch with similar size (DBH), belonging to the same plant functional type (PFT), and characterized by a dynamic plant density (). Patches and cohorts are spatially implicit; that is, they do not have explicit spatial positions (Medvigy et al. 2009).

Previous studies have demonstrated the ability of the ED2.2 model to reproduce important aspects of carbon and water dynamics in tropical ecosystems (Longo et al. 2019b). In particular, it was shown that ED2 could correctly simulate reductions in above-ground biomass of Amazon forests subjected to drought experiments (Powell et al. 2013), capture mortality rates and above-ground biomass stocks on Barro Colorado Island (BCI), Panama (Powell et al. 2017), and represent leaf and biomass spatial and temporal variability in tropical dry forests (Xu et al. 2016). Recently, a new plant functional type accounting for the lianescent growth form was implemented in the ED2 model by di Porcia e Brugnera et al. (2019). The liana PFT was extensively calibrated by Meunier et al. (2020), but that calibration did not include radiative transfer parameters.

### The radiative transfer model of ED2

In ED2, the radiative transfer is modelled as a multi-layer version of the two-stream model (Liou 2002; Sellers 1985) applied to three broad spectral bands: visible (400-700 nm), solar (near and shortwave) infrared (700-3000 nm) and thermal infrared radiation (3-15 µm). The two-stream approach is the core canopy radiative transfer model of multiple global vegetation models, including JULES (Best et al. 2011) and CLM (Lawrence et al. 2019). We focused on the visible and solar infrared bands in this study.

In ED2, the radiation regime is based on single scattering and backscattering coefficients computed from prescribed PFT-specific leaf transmissivity (and for the visible and solar infrared bands, respectively) and reflectivity (and) coefficients, as well as a leaf orientation parameter and the vertical structure of the canopy. Leaf orientation is a PFT-specific parameter that determines the average leaf surface area in the direction of the radiation beam. It varies between -1 (all leaves are vertically oriented) and +1 (all leaves are horizontally oriented), with 0 meaning randomly oriented leaves. The vertical structure of the canopy is determined by the size distribution of the plant cohorts within each patch. Each cohort is represented by a flat-topped layer whose vertical position is scaled with its diameter *DBH*_*k*_ according to a Weibull function :

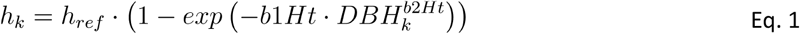

where *h*_*ref*_, *b*1*Ht* and *b*2*Ht* are PFT-specific allometric parameters. As opposed to trees, liana vertical position is not only determined by their size but also by the height distribution of the trees in the vicinity: in the model, lianas can only only overtop trees belonging to the same patch by a small threshold because they lack self-supporting tissues (Meunier et al. 2020). Following Kurzel et al. (2006), liana initial height was determined by their size and the height distribution of the surrounding trees so that all lianas with a stem diameter > 3cm reached the top of the forest canopy, resulting in a liana vertical clumping.

The optical thickness of each layer is computed from the total plant area index which is the sum of the wood and the clumping-corrected leaf area index. In this study, we chose to neglect the wood area given the low number of observations of liana wood optical properties, its limited impact on the radiative transfer, as well as to limit the number of parameters to calibrate. Therefore the total plant area index of each cohort (Ф_*K*_) was given solely by the leaf area index (*LAI*_*k*_). In ED2, each cohort crown occupies the full patch horizontal area (i.e. cohort leaves are distributed across the entire patch) but plant leaf area is corrected by a PFT-specific clumping factor (Ω). Ω is a purely horizontal clumping factor reducing the cohort light interception. In ED2, it is a constant that does not depend on the direction of the incident beam but is intended to compensate for the fact that radiative transfer is solved in 2D rather than in 3D. Ωcan vary between 0 and 1 and these numbers respectively represent the two extreme situations of a perfectly clumped (infinite LAI over a infinitesimally small area and null effective light interception, Ω=0) and an evenly spread canopy (which captures the maximum amount of light per unit of leaf area, Ω=1). Cohort LAI (*LAI*_*k*_) is the product of the plant-level leaf biomass (*B*_*leaf,k*_ which scales with diameter through another allometric equation), the plant density (*n*_*k*_) and the PFT-specific dry matter content (*LM A*). Therefore, the cohort plant effective area index Φ_*k*_ is given by:

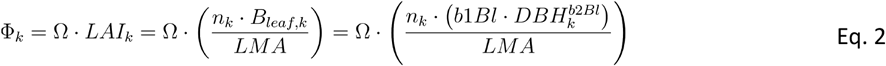

with *b*1*Bl* and *b*2*Bl* the PFT-specific leaf biomass allometric intercept and slope coefficients, respectively. The relevant parameters are again further described in Table 1. The complete description of the radiative transfer model of ED2 can be found in Appendix A, in Longo et al. (2019a), or in Shiklomanov et al. (2020).

Recent work (Shiklomanov et al. 2020) has refined the ED2 model’s visible and solar infrared spectral resolution to nanometer-scale bands by coupling it with PROSPECT-5, which provides fine leaf spectral properties, so as to allow direct comparisons to multi- and hyperspectral data. This model version (a.k.a. ED-RTM) is instantaneous (the model runs for a single time step at user prescribed time and date) and allows computing radiative transfers within the canopy and the forest albedo down to the nanometer spectral resolution.

## Experimental workflow

### Literature meta-analysis

We first collected published spectral data through an extensive literature search on science research engines (Web of Science and Google Scholar) with a combination of the following key-words: “liana/woody vine”, “spectrum”, “reflectance” and “canopy/leaf/leaves”. We compiled records presenting leaf spectral measurements of both tropical lianas and trees, or canopy spectral measurements covering different levels of liana infestation in tropical forests. In the latter case, we only analyzed the measurements with the lowest and highest liana coverages (hereafter referred to as liana-free and liana-infested stands) to capture the maximal differences in canopy spectra generated by liana infestation. We focused on data that overlapped the spectral region of interest: the visible and the solar infrared regions of the spectrum (400-2500 nm). Seven references emerged from our extensive search. These references could be categorized in three groups: studies that investigated leaf-level reflectance only (3), patch-level reflectance only (3), or both (1), see Table 2. In total, we compiled 1689 leaf spectra (1162 from lianas, 527 from trees) and eight patch-level average canopy spectra. All five study sites were located in Latin America (Panama, Costa Rica and Bolivia).

**Table 2:** Basic information related to the studies included in the meta-analysis. FTS = Fort Sherman, PNM = Parque Natural Metropolitano, SRPN = Santa Rosa National Park, NKMNP = Noel Kempff Mercado National Park.

| Reference | Study site(s) [Short name] | Leaf reflectance spectrum range (nm) | Canopy reflectance spectrum range (nm) | Data source | Date of collection | Number of spectra<sup>1</sup> |
| --- | --- | --- | --- | --- | --- | --- |
| Castro-Esau et al. (2004) | FTS, Panama [Castro, FTS]<br>PNM, Panama [Castro, PNM] | 450-950 | - | Digitized | 2013/03<br>(dry season) | 1 (L), 1 (T) |
| Guzmán et al. (2018) | SRNP, Costa Rica [Guzmán] | 450-950 | - | Original | 2017/05-07<br>(wet season) | 14 (L), 21 (T) |
| Sanchez et al. (2009) | FTS, Panama [Sanchez, FTS]<br>PNM, Panama [Sanchez, PNM] | 400-2500 | - | Original | 2004/08<br>(wet season) | 1146 (L), 504 (T) |
| Kalacska et al. (2007) | PNM, Panama [Kalacska] | 400-2500 | 450-2250 | Digitized | 2004/12<br>(dry season) | 1 (L), 1 (T)<br>1 (LF), 1 (LI) |
| Marvin et al. (2016) | Gigante, Panama [Marvin] | - | 400-2500 | Digitized | 2012/02<br>(dry season) | 1 (LF), 1 (LI) |
| Foster et al. (2008) | NKMNP, Bolivia [Foster] |  | 400-2500 | Digitized | 2003/04<br>(dry season) | 1 (LF), 1 (LI) |
| Sanchez et al. (2006) | PNM, Panama [Sanchez] | - | 400-2400 | Digitized | 2012/07<br>(wet season) | 1 (LF), 1 (LI) |

<sup>1</sup>Leaf-level studies: L = Liana, T = Tree
Stand-level studies: LF = Liana-free patch, LI = Liana-infested patch.

Only two of the studies, Guzmán et al. (2018) and Sanchez-Azofeifa et al. (2009), made their raw data available. For those studies that did not provide their data, we digitized the spectra using the software Plot digitizer (v.2.6.8, http://plotdigitizer.sourceforge.net/) from the relevant figures. At the leaf-level, two studies (out of the 4) reported spectral data only for parts of the visible and the near infrared (450-950 nm). All four studies that investigated patch-level reflectance differences with liana infestation included a large fraction of the visible and the near/short-wave infrared (Table 2). The significance of the liana impact was evaluated by testing whether the 95% confidence intervals of the difference of the spectra (liana *vs* tree, liana-free *vs* liana-infested) in a specific region included zero.

### Model calibration

To calibrate the leaf and the canopy spectra, we used the R package PEcAnRTM (https://github.com/PecanProject/pecan/tree/develop/modules/rtm). This approach uses a Bayesian framework that provides the joint probability posterior distribution of the parameter set, thus capturing both the uncertainties and correlations among parameters. The independent prior distributions used in this study are included in Table 1. All but the priors of the leaf biomass allometric coefficients are uninformative. The residual variance*σ*^*2*^ was assigned an uninformative inverse gamma prior as in Shiklomanov et al. (2016). We fit the respective models using the Differential Evolution with Snooker Update (“DEzs”) Markov-Chain Monte Carlo (MCMC) sampling algorithm (ter Braak and Vrugt 2008) as implemented in the R package BayesianTools (Hartig et al. 2019). For each inversion, we ran five independent chains until convergence with a burn-in of 10,000 and 100,000 iterations for leaf and canopy scales, respectively. We evaluated the quality of the model calibration at the leaf- and the stand-levels by fitting modelled and observed reflectance values and by computing several error statistical metrics, see Appendix B for more details.

### Estimating leaf optical traits from leaf spectra

For all studies for which we had leaf-level reflectance data, we estimated all five PROSPECT-5 leaf-level parameters (Figure 1, step 1), through the exact same spectral inversion procedure described by Shiklomanov et al. (2016). We repeated the PROSPECT-5 inversion for each site/study and each PFT. When raw data were available, we fitted PROSPECT-5 parameters individually to each plant species. After calibration, we aggregated all study/site posteriors to the PFT-level (liana or tree) by attributing a similar weight to each of them.

### Refining leaf optical trait estimates from canopy spectra

For the studies for which patch-level reflectance data were available, we followed Shiklomanov et al. (2020) to carry out a similar calibration scheme using ED-RTM. The calibrated model parameters include the leaf parameters of PROSPECT-5 plus the leaf allometric coefficients, the leaf orientation and the clumping factor (Figure 1, step 2). We used the aggregated posterior distributions from step 1 as informative prior distributions for the PROSPECT-5 parameters. The priors of leaf biomass allocation coefficients were informed by an independent literature review (Gehring et al. 2004; Gerwing and Farias 2000; Putz 1983; Smith-Martin et al. 2019; Falster et al. 2015; Schnitzer et al. 2006) through which we collected all leaf biomass allometric data comparing trees and lianas (Meunier et al. 2020). Informed priors of leaf biomass allometric coefficients were built based on the fit of the allometric equation of ED2 to these data using the PEcAn.allometry package (https://github.com/pecanproject/pecan/tree/develop/modules/allometry).

In ED-RTM, we simulated two competing PFTs, respectively accounting for lianas and tropical trees. For the purpose of the canopy inversion, we assumed that all trees at a given site shared the same spectral property distributions, and all lianas likewise shared another set of spectral property distributions. As for the leaf-level studies, we performed a separate calibration for every single study and site (Table 2) and aggregated the posterior distributions giving the same weight to every calibration. This approach allowed a better fit for every leaf canopy spectra while accounting for the inter-site variability of liana and tree properties.

When both leaf- and patch-level reflectance data were available (Kalacska et al., 2007), we performed a similar two-step calibration. Yet as compared to the other stand-level calibrations, we only used the site-specific PROSPECT-5 posterior distributions as priors in the second step to derive canopy structural traits. To improve the vertical resolution of the radiative transfer, we prevented cohort fusion during the calibration of ED-RTM. Therefore every single plant cohort (liana or tree) consisted of a single individual (see Supplementary Figure F5).

### Evaluating the impacts of liana-specific traits

To determine the impact of lianas on forest radiative transfers and biogeochemical cycles, posterior distributions of leaf and canopy traits of both PFTs resulting from the two-step calibrations were used to parameterize the full version of the ED2.2 vegetation model (Figure 1, step 3). We generated two ensembles of 100 simulations each in which liana radiative traits were sampled either from the liana (“liana” runs) or the tree (“reference” runs) posterior distributions. Liana leaf biomass allometric parameters (*b*1*Bl* and *b*2*Bl*), dry matter (*LM A*), and all tree parameters were systematically sampled from their proper PFT distributions. This allowed us to evaluate the impact of the leaf radiative parameters (the visible and infrared transmissivity and reflectivity coefficients - *τ*_*vis*_, *ρ*_*vis*_, *τ*_*IR*_, and *ρ*_*IR*_ - leaf angle ω and leaf clumping Ω) on forest functioning while conserving identical ecosystem LAI.

We ran the model on the 50-ha plot of BCI, Panama, which is an old-growth seasonally moist lowland tropical forest. Previous ED2 simulations demonstrated the model’s capacity to reproduce land surface fluxes (di Porcia e Brugnera et al., 2019) and several features of the liana-tree competition there (Meunier et al. 2020). BCI is characterized by an average (years 2003-2016) annual rainfall of about 2640 mm (Detto et al. 2018) and a well-marked 4-month dry season (total rainfall between late-December and mid-April is 175 mm on average). We used the only available liana inventory (2007) and the tree inventory that was carried out right after (2010) to prescribe the vegetation initial conditions. These censuses include all individuals whose DBH is larger than 1 cm in the 500 m x 1000 m plot (Condit et al. 2019; Schnitzer et al. 2012). The 50-ha site was divided into an initial number of 1250 patches in a regular grid of 20 x 20 m. Initial liana density averaged 1429 individuals per hectare and varied significantly from liana-free to liana-infested patches (liana density standard deviation was 1013 individuals per hectare). Model simulations were run for five years starting in 2007 using the meteorological drivers locally available (Powell et al. 2017; Powell et al. 2018). We also used the observed carbon and energy land fluxes obtained with the eddy-covariance method to benchmark the modelled productivity, evapotranspiration, and albedo (Pau et al. 2018). The other model parameters for the liana and the tree PFTs are exhaustively described in Meunier et al. (2020) and in Longo et al. (2019a).

We analyzed the differences between model outputs with a special focus on the energy budget and the carbon cycle. These differences were evaluated as both absolute (“liana” runs minus “reference” runs) and relative (absolute differences normalized by the “reference” runs) changes.

In addition, we also simulated the instantaneous radiative transfers at the same site using ED-RTM and finer patch and cohort resolutions (the regular grid of 20 x 20 m kept unfused and every single plant as a ‘cohort’) and the posterior median of every single parameter. Again, liana optical traits were either assigned liana- or tree-specific values. We then related the changes of forest albedo and understorey light (defined as the downward radiation below the shortest plant or the radiation reaching the ground) to the degree of liana infestation by fitting the coefficients *y*_*max*_ and *b* of the following equation using the ‘nlsLM’ function of the minpack.lm R package (Elzhov et al. 2016):

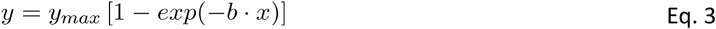

where *y* is the response (forest albedo in the visible and in the infrared, or the understorey light) and *x* is the contribution of lianas to the ecosystem LAI.

Finally, we ran uncertainty analyses of every single model used in this study (PROSPECT-5, ED-RTM, and ED2.2) with PEcAn (LeBauer et al. 2013). In short, PEcAn combines the uncertainty of model parameters after calibration with a univariate sensitivity analysis to estimate the contribution of each parameter to the overall predictive uncertainty, see Appendix C for details. All intervals presented in this study are 95% confidence intervals (CI).

### Independent evaluation of model outputs

We validated the main model outputs using independent datasets. In particular, we evaluated the impacts of lianas on forest albedo using WorldView-3 images and on light penetration through the canopy using GatorEye UAV-LiDAR.

#### WorldView-3

To validate the changes in albedo due to liana infestation, we obtained a WorldView-3 image of central Panama acquired on March 24, 2016. The image has a 2 m spatial resolution, 2.6 degree mean off-nadir view, 67.9 degree mean sun elevation, with 11.5% cloud cover. The image was corrected to surface reflectance using Fast Line-of-sight Atmospheric Analysis (Cooley et al. 2002; http://www.harrisgeospatial.com). The scene included a cloud-free area of georeferenced tree crowns on Gigante Peninsula (south of BCI) where liana infestation of 544 trees was tallied from the ground with binoculars using the liana crown occupancy index (COI, see Marvin et al. 2016). This resulted in over 8000 pixels with ground referenced infestation scores. To investigate the effect of liana infestation on forest albedo, we compared the reflectance of crowns with extreme levels of liana occupancy (0.05 < COI < 0.25 *vs* COI > 0.75) across six spectral bands (from blue to NIR-2), using independent nonparametric Mann–Whitney U tests for each band and the COI category as treatment. The two liana-infestation levels resulted in large numbers of samples in both groups (5866 and 387 for low and high COI, respectively totalling 2.35 and 0.15 ha).

### UAV LiDAR

To evaluate whether liana-rich stands differed in their light penetration, GatorEye UAV-borne LiDAR data obtained on BCI in February 2019 was combined with a local dataset of COI. Tree crowns were mapped for a total of over 2133 individuals. Each crown was manually delineated from UAV derived orthomosaic of the 50-ha plot of BCI, as described in Park et al. (2019). Each delineated tree crown was then visited in the field and a liana crown occupancy index was recorded from the ground. UAV-borne LiDAR data was collected at 905 nm, using the GatorEye Unmanned Flying Laboratory (see www.GatorEye.org for detailed description and data access) at the 50-ha plot on Barro Colorado Island. GatorEye Gen 1 uses a DJI Matrice 600 Pro hexacopter, equipped with a Velodyne VLP-16 puck lite sensor with manufacturer calibrated intensity, as well as hyperspectral and visual sensors not used in this study. Flights were conducted at 55 m AGL, with an 80% sidelap, and at an average speed of 40 km/h. The LiDAR spatial footprint has a diameter of 3-9 cm at 10-30 meters distance, resulting in an average canopy height (30 m AGL) footprint of 7.5 cm diameter. Here we compare the manufacturer calibrated intensity as measured by the LiDAR. The intensity varies with reflectivity and distance. Yet, as the distance from sensor to tree canopies was approximately the same across the entire plot and we average thousands of returns per quadrant, intensity is expected to provide a good estimation of relative differences in reflectivity and leaf density among the sample areas. In other words, LiDAR intensity should differ between liana-free and liana-infested tree crowns if the projected leaf area and traits within the beam footprint differs - which could be caused by differences in clumping, leaf angles, or leaf reflectivity. We compared the relative (scaled between 0 and 1) LiDAR intensity of crowns with extreme levels of liana occupancy (COI < 0.25 *vs* COI > 0.75), using a nonparametric Mann–Whitney U test with the liana infestation level as treatment. These two categories resulted in a large number of individual crowns in both groups (1486 and 417 for low and high COI, respectively).

## Results

### Leaf biochemical and canopy structural trait differences between growth forms

Most collected leaf-level studies agreed on the direction of the discrepancies between liana and tree leaf spectra (Appendix D). Five out of the six leaf-level collected references indicated that liana leaves were significantly more reflective than tree leaves in the visible (especially in the green peak) until the red edge (680-700 nm). The sixth study, Sanchez (FTS), showed no differences, on average, between liana and tree leaves in the visible region of the spectrum. In the near infrared (700-1400 nm), all studies agreed on a significantly lower reflectance of liana leaves at least until 950 nm (the upper limit of spectral measurements in Castro (FTS), Castro (PNM) and Guzmán, see Table 2). In the shortwave infrared (1500-2500 nm), liana leaves were variably characterized as having on average higher (Kalacska), lower (Sanchez, PNM) or the same (Sanchez, FTS) reflectance as tree leaves. At the canopy level, all four studies found higher reflectances for liana-infested patches in the visible and short-wave infrared, and all but Kalacska also found higher reflectance in the near-infrared (Figure C2 and Table C1).

Both PROSPECT-5 and ED-RTM were able to accurately reproduce observed leaf and canopy spectra from individual studies across the whole spectrum (Appendix E). At the leaf level, r^2^ of observed *vs* modeled leaf reflectance values grouped by wavelength reached on average 0.91, with no values lower than 0.75 (Supplementary Figure E3). Overall simulated leaf spectra reproduced discrepancies between liana and tree leaf reflectance values both in the visible and in the infrared (Supplementary Table D1). Similarly, at the canopy level, model performance after model calibration was good with a mean r^2^ of 0.93 for observed *vs* simulated reflectance values across all wavelengths, and none lower than 0.71 (Supplementary Figure E3). Observations of reflectance discrepancies between forest stands with different infestation levels were also well reproduced by the best set of parameters, both in the visible and in the infrared (Supplementary Table E1).

Liana and tree parameter distributions had large variances and therefore largely overlapped across and within studies. However, we did observe significant differences in central tendencies. All calibrations predicted lower chlorophyll contents in liana leaves as compared to tropical tree leaves (Supplementary Figure F1). The difference between chlorophyll content in liana and tree leaves reached -11.1 µg cm^-2^ when averaging all studies (Table 1). Similarly, all studies/sites parameter calibrations predicted lower carotenoid contents in tropical liana leaves (on average -4.7 µg cm^-2^) and a smaller number of stacked layers (−0.3) corresponding to thinner leaves in liana species. The calibrations predicted either no differences or smaller dry matter content in liana leaves (*LMA* decreased on average by -0.022 kg m^-2^ for liana leaves). In addition, significant differences of water layer thickness *C*_*w*_ emerged in all studies/sites but one (Castro, PNM), leading to a mean water content increase of +3.7% in liana leaves (once combined with *LMA*). All together, those traits resulted in significant increases in liana leaf transmittance (+ 0.02 in the visible and + 0.01 in the solar infrared) and reflectance in the visible (+ 0.004), as well as a significant decline in liana leaf reflectance in the near-infrared (−0.01), see Figure 2A (as well as Supplementary Figure F2).

**Figure 2:**
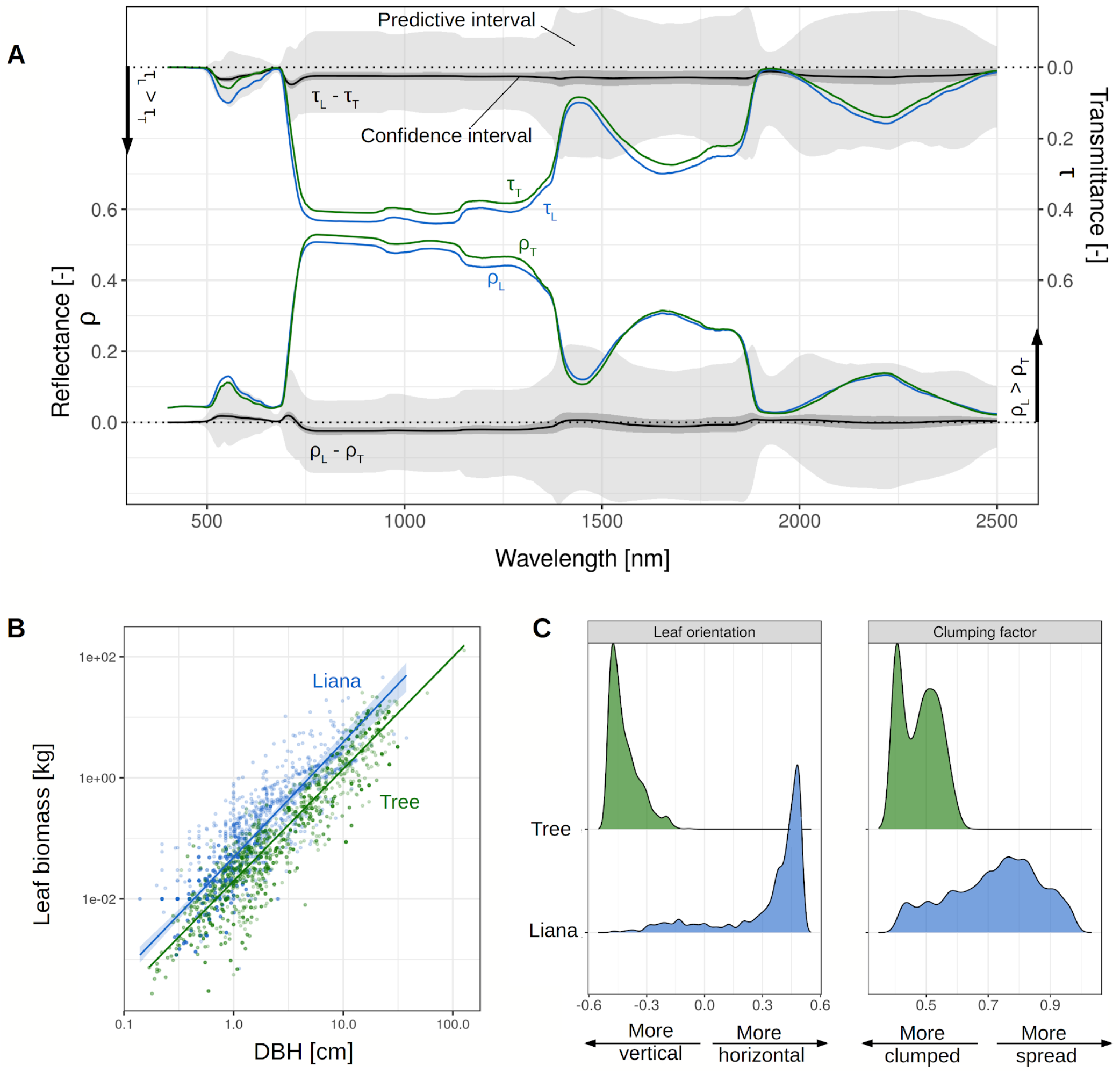
Liana (blue) and tree (green) optical and canopy structural parameters, resulting from the leaf and canopy spectral calibrations. In A, liana and tree mean reflectance (ρ_L_ and ρ_T_, respectively) and transmittance (τ_L_ and τ_T_, respectively) are plotted together with their differences (ρ_L_ - ρ_T_ and τ_L_ - τ_T_) at the nanometer resolution. The light and dark grey envelopes respectively represent the 95% predictive and confidence intervals of the differences (liana - tree) resulting from 500 liana and tree PROSPECT-5 simulations sampled from the growth-form specific posterior distributions. In B, liana and tree mean (solid lines) allometric allocations to leaf biomass together with their confidence intervals (shaded envelopes) are superimposed on the data that served to constrain the prior distributions (collected through an independent meta-analysis). In C, we compare the posterior distribution densities of liana and tree leaf orientation (left) and clumping factor (right), resulting from the calibration of the canopy spectra.

Leaf allometric intercepts already differed between growth forms according to the literature meta-analysis (Table 1) and were further discriminated by model calibration (Supplementary Figure F1). The slope of the leaf biomass allometric equation was slightly larger for lianas, before and after calibration (Table 1 and supplementary Figure E1). Together, these leaf biomass allometric coefficients drew the picture of a larger carbon allocation to leaf biomass by lianas than by trees of similar DBH across all sizes (Figure 2B). All four studies that measured the impact of liana coverage on canopy reflectance revealed, after calibration, more horizontal leaves for lianas than for trees (Supplementary Figure F1), resulting in flatter leaf angles for lianas (Figure 2C). In addition, lianas were systematically predicted to have larger clumping factors (*i.e*. larger light interception per unit of leaf area) as compared to trees (Figure 2C and Table 2).

### Impact of liana traits on forest biogeochemical cycles

The leaf biochemical and canopy structural traits of lianas deeply impacted the energy balance and the carbon cycle of the forest, as simulated by ED2 (Figure 3). Liana leaf traits increased in forest albedo: over the five years of simulation, the energy reflected back to the atmosphere increased on average by 17.1% in the PAR mainly because of the higher reflectivity of liana leaves in the visible region and by 11.7% in the infrared driven by more horizontal angle distribution of liana leaves and larger clumping factors (Appendix C). This corresponded to an additional 3.6 W m^-2^ of energy that was sent back to the atmosphere (Supplementary Figure F3). A Larger fraction of the penetrating forest solar radiation was absorbed by the canopy in the runs where liana-specific traits were attributed to lianas (*i.e*. “liana” runs), with an average increase by 7.4% and 7.8%, respectively in the visible and the solar infrared (or + 8.2 W m^-2^ in total). Hence, fewer photons reached the ground resulting in darker understories (−11.8 W m^-2^ as summed up over the visible and the infrared, corresponding to a 22% decrease) and slightly cooler soils (on average -0.5°C in the topsoil over the five year of simulations).

**Figure 3:**
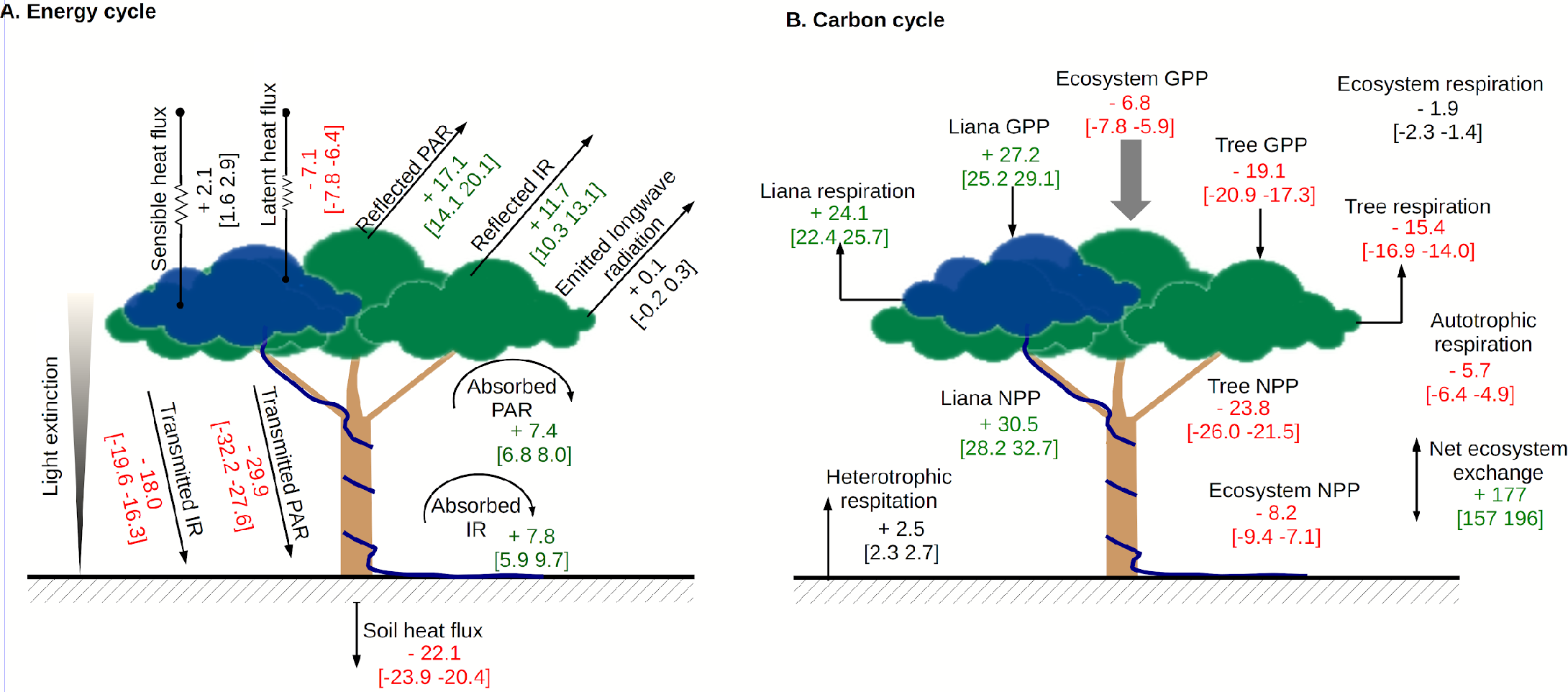
Mean relative changes (%) together with their confident intervals of the energy (A) and carbon (B) cycle fluxes resulting from the implementation of the liana radiative model parameters. These changes are relative to the fluxes simulated when lianas were assumed to have the same radiative and structural parameters as trees. Fluxes are coloured in red (respectively green) when the mean relative changes of the corresponding fluxes are lower than -5% (respectively higher than +5%).

The ecosystem-level increase of albedo and decrease of understorey light were mainly driven by the liana-rich patches on BCI (Figure 4). In liana-infested patches, the increase of albedo reached up to 0.05 in the infrared and 0.002 in the visible, while light reaching the ground could be reduced by more than 50%. On BCI, the liana optical traits increased the shortwave albedo by + 0.015 on average, which improved the fitness of the simulated outgoing shortwave radiation when compared to the fluxtower data: RMSE of the modelled *vs* observed reflected shortwave radiation decreased from 2.2 to 1.7 W m^-2^ once liana traits were accounted for (Supplementary Figure F4).

**Figure 4:**
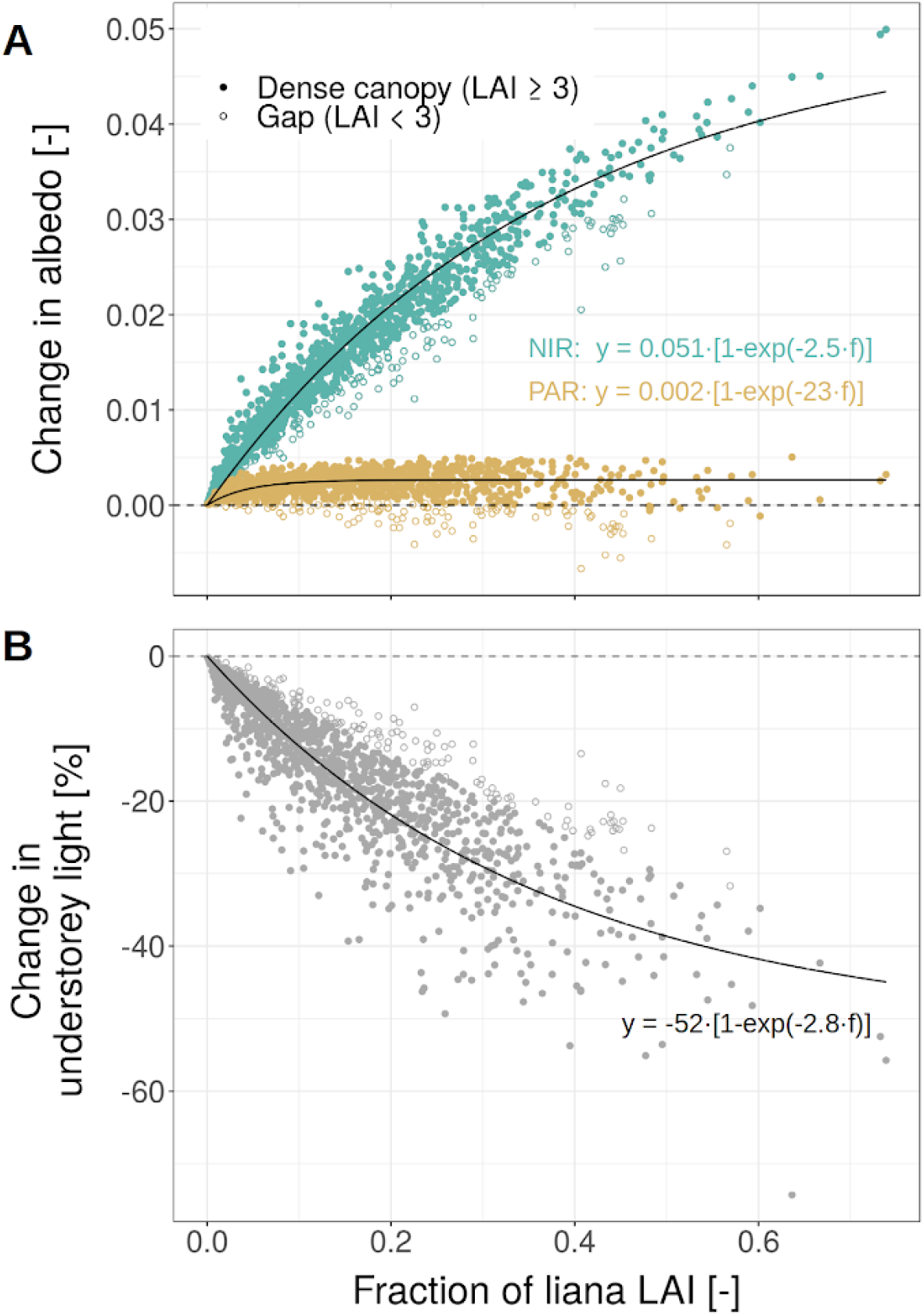
Changes in albedo (A) and understorey light availability (B) as a function of the liana infestation, expressed here as the contribution of lianas to the ecosystem LAI. In A, the impact was split into changes of PAR (yellow) and infrared (green) albedo, and in both subplots we distinguished dense vegetation patches (LAI ≥ 3, closed symbols) and gaps (LAI < 3, open symbols). Fits were applied to all data points (dense canopies and gaps together).

The larger values of the leaf orientation parameter for lianas (*i.e*. more horizontal leaf distribution) and clumping parameters (*i.e*. more evenly distributed leaves) made lianas very efficient at light interception (intercepted PAR by lianas increased by 48% on average when liana optical traits were accounted for). This translated into a large increase in both gross and net productivity of the liana PFT (+27.2% and +30.5%, respectively) while trees were negatively impacted by those liana-specific traits (tree GPP and NPP decreased by 19.1% and 23.8%, respectively). Tree productivity decline was not compensated by liana gains: compared to the reference runs, ecosystem gross and net productivity decreased in the “liana” runs by 6.8% and 8.2%, respectively. Combined with a slight increase of heterotrophic respiration, the net ecosystem productivity was reduced by 1.6 T_C_ ha^-1^ yr^-1^ when liana-specific optical traits were accounted for. After 5 years of simulation, ecosystem AGB increased more slowly in the “liana” runs as compared to the “reference” ones (reduction of 4 T_C_ ha^-1^): tree AGB increase was reduced by 4.5 T_C_ ha^-1^ while liana AGB was enhanced by 0.5 T_C_ ha^-1^ (+11%). The simulated seasonal cycle of latent heat and gross primary productivity were both slightly improved when liana-specific optical traits were taken into account (Supplementary Figure F4).

### Independent evaluation of model outputs

WorldView-3 images confirmed that liana-infested canopies were characterized by significantly higher albedos in the near-infrared (+ 0.051 on average, p-value < 2e-16) and in the green peak (+ 0.002, p-value = 0.004), see Figure 5A. These numbers matched the asymptotes predicted by the radiative transfer model under extreme levels of liana infestation (Figure 4).

**Figure 5:**
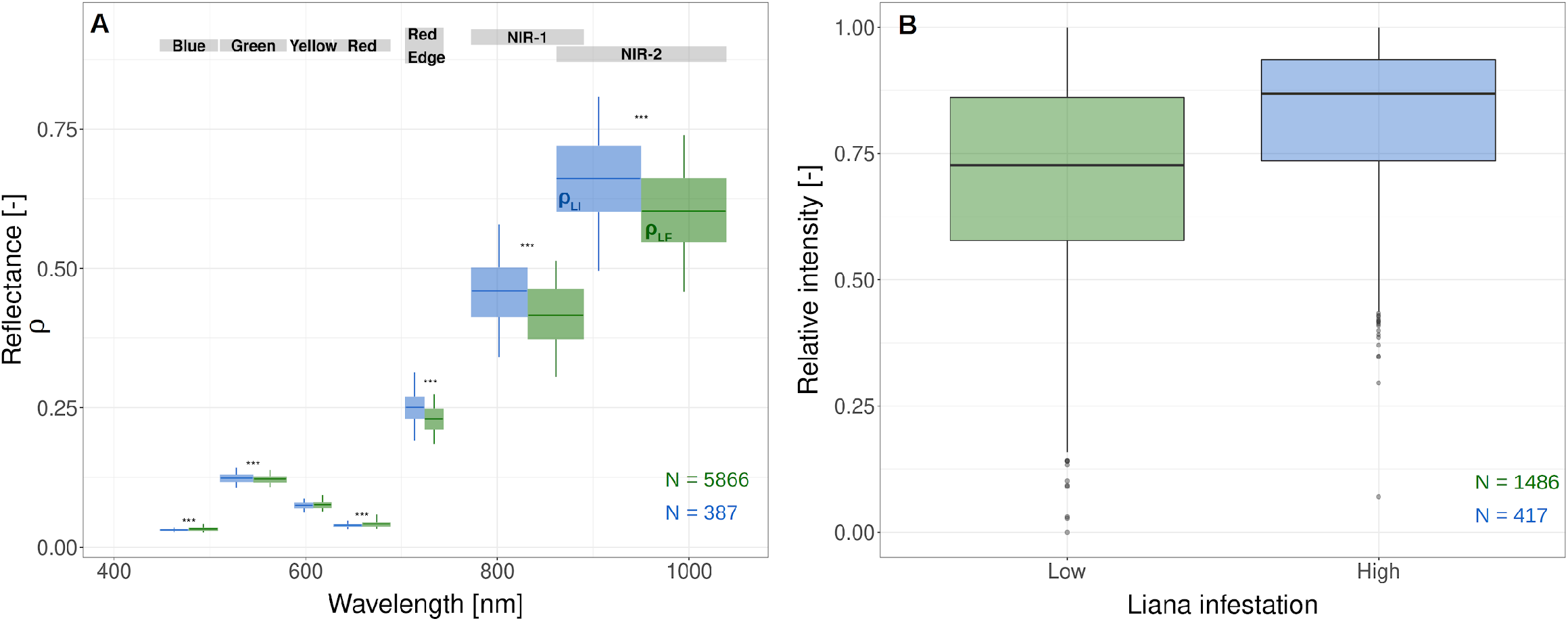
Independent evaluation of the liana impacts on forest albedo (A) and light penetration (B). In A, we compare the mean reflectance of liana-free (green, ρ_LF_) and liana-infested (blue, ρ_LI_) canopies. Boxplot widths correspond to half the size of the bands in the Worldview-3 images (indicated above). In B, the relative intensity detected by UAV-borne LiDAR measurements is compared between low and high liana infestation. Higher values indicate higher return rates (smaller light penetration into the canopy).

We could not validate individually the differences in canopy clumping and leaf angle of liana species but the GatorEye UAV-LiDAR confirmed the more important canopy closure of liana-rich crowns resulting from the combination of such features (Figure 5B). Larger COI were associated with greater LiDAR intensities (p-value < 2e-16), which indicates that there is a greater and more reflective leaf area within the footprint of the laser. As liana leaves tend to have lower reflectance around 900 nm (Figure 2A), the greater return intensity of infested crowns is likely caused by a higher LAI of more horizontal leaves, more evenly spread canopies, or both, which is consistent with the posterior distribution discrepancies between growth forms (Figure 2C and Table 1).

## Discussion

### Ecosystem-level impacts of lianas

This study is the first to comprehensively investigate the role of lianas on the energy budget and radiative transfer of tropical forests. According to our model simulations, lianas profoundly alter the forest energy cycle (Figure 3A). As hypothesized, liana optical traits are responsible for an increase in albedo and a decrease in light penetration through the canopy, with the most dramatic impacts in heavily infested forest patches (Figure 4). By increasing the light competition, liana traits also severely reduce tree carbon assimilation and the positive shift in liana productivity does not compensate for the tree drop, resulting in an overall decline of the ecosystem productivity (Figure 3B). Since liana abundance is increasing in the Neotropics (Phillips 2002; Schnitzer and Bongers 2011), these findings might translate into dramatic changes in the functioning of those tropical forests. At a large scale, forest darkening and exacerbated light competition could translate into a global weakening of forest carbon sink strength, not only reducing forest growth rates and storage capacity but also completely disrupting the competitive hierarchies among tree species, as observed in temperate forests (Blondeel et al. 2020; De Lombaerde et al. 2020).

Yet, we note that these negative impacts could be mediated or even counterbalanced by other liana-induced effects. By reducing the solar radiation reaching the ecosystem (especially in the infrared), overtopping lianas could act as a protective mantle isolating tree leaves from rising temperatures and the resulting exacerbated evaporative demand expected in the tropics (Konapala et al. 2020). Lianas could also help maintain more appropriate soil conditions (temperatures and water content) and hence protect soil carbon reservoirs as well as the critical biota living therein, which are currently threatened by increasing soil temperatures (Mitchard 2018). Finally, as degraded forests with more open canopy (and hence warmer understorey temperatures) are more flammable during mild/moderate droughts (Longo et al. 2020), the susceptibility to fires could also be reduced in liana-infested patches.

Finally, our results suggest that the increase in liana abundance observed in the Neotropics (Phillips 2002; Schnitzer and Bongers 2011) could be driven by the higher efficiency of lianas to intercept light combined with an overall decrease in cloudiness in the area (Arias et al. 2011; Fu et al. 2013). The liana-specific leaf and canopy traits identified in this study coupled to earlier observations of efficient hydraulic architectures (van der Sande et al. 2013; Chen et al. 2015; Chen et al. 2017) could indeed allow lianas to thrive through light-rich extended dry seasons and more frequent drought stress episodes, during which lianas have been shown to grow more than co-occurring trees (Schnitzer 2005; Schnitzer and van der Heijden 2019).

### Study limitations

The most important limitation of this study is the representativity of the data that we collected through our meta-analysis. Even though we were able to compile a large number of liana leaf spectra (>1000), they all originated from Latin America (mostly from Panama). The stand-level spectra that we collected also all came from the Neotropics (again mostly from Panama) and their number was more limited as we could not obtain any raw data. The extrapolation of the impact of lianas on all tropical rainforests globally should therefore be performed with caution until additional liana spectra with more diverse origins have been collected and assimilated into the model. In particular, leaf sampling in regions where lianas do not seem to be increasing (Bongers et al. 2020) could lead to a breakthrough for unraveling the mechanisms driving liana proliferation. In addition, a publication bias could not be ruled out in our meta-analysis as the number of collected studies was too small to calculate publication bias diagnostics such as funnel plots (Koricheva et al., 2013). Yet, the inversion of the leaf spectra led to a pattern (lianas grow cheaper, thinner leaves, with lower pigment concentration and lower leaf mass per area), which is consistent with multiple local observations (Castro-Esau et al 2004; Sánchez-Azofeifa et al. 2009), the pan-tropical analysis of Asner and Martin (2012) who found smaller and no differences or lower area-based light capture-growth chemical concentrations in liana leaves, and higher leaf turnover rates in liana-rich plots (van der Heijden et al. 2015).

Furthermore, the ED2 (and ED-RTM) representation of infinitely thin crowns covering the entire patch and overtopping lianas probably contributes substantially to the large impact of lianas on forest albedo, and energy balance. These model structural uncertainties, even if they have been shown to be less critical than model parameter uncertainties for ED2 (Shiklomanov et al. 2020) and are compensated in the model and in our analyses (clumping factor, cohort size limit), need to be accounted for in the future, by investigating the impact of plant crown size (Dietze et al. 2008) and crown depth representation (Fisher et al. 2018; Fisher et al. 2010) in the canopy radiation model. Yet, the effects of liana abundance on albedo predicted by the model are consistent with those observed in the original stand-level spectral data, as well as the independent Worldview-3 multispectral and GatorEye UAV-LiDAR data, suggesting that the impacts of any structural biases are either minor or related to compensating errors that are not apparent in any of our diagnostics. Additional independent field data on the leaf angle distribution and leaf clumping would help rule out such compensating errors, but it is worth noting that the current parameter estimates are, at least qualitatively, consistent with our understanding of liana canopy morphology and liana ecology, more broadly. Our results are indeed in agreement with experimental observations of a steep increase in forest net productivity once lianas are removed (van der Heijden et al. 2015) and further suggest that light competition is the critical driver of competition, as expected in dense tropical ecosystems (Bongers and Sterck 1998; Poorter et al. 2003). The strong reduction in light penetration induced by liana infestation was also evidenced in several liana removal experiments (Estrada-Villegas and Schnitzer 2018).

### Perspectives

Our study opens new avenues to estimate the contribution of lianas on seasonal and interannual changes of forest albedo observed at large scales (Ahlström et al. 2017; Asner and Alencar 2010; Brienen et al. 2015). Since liana leaf biomass, relative to trees, can increase during the dry season (Schnitzer 2005; Avalos and Mulkey 1999) and is expected to soar on the longer term (Schnitzer and Bongers 2011; Phillips 2002), liana-induced shifts in albedo could be visible both at short- and long-timescales. Vegetation model simulations could help estimate the future impact of lianas on tropical forest albedo and functioning, more broadly under contrasting global scenarios.

Our results (Figure 4 in particular) also suggest that multispectral sensors (alone or in combination with thermal cameras (Guzmán et al. 2018; Yuan et al. 2019) or LiDAR) should be able to detect forest stands characterized by high liana coverage. In a related study, Visser et al. (2021) show how radiative transfer models can assist in the estimation of liana traits from hyperspectral images of liana-infested canopies just as it is currently achieved for trees (Gong et al. 2003; Meroni et al. 2004; Serbin et al. 2015). This is because liana-rich patches are similarly sensitive to liana parameters, just like liana-free forest stands are to tropical tree parameters (Appendix C).

This study is a first step toward a comprehensive understanding and quantification of the impact of lianas on tropical forests. The current belief is that the increase in liana abundance driven by climate change and exacerbated by anthropogenic pressures could accelerate global warming by decreasing the carbon sequestration capacity of ecosystems (Reis et al. 2020; Schnitzer and Bongers 2011; van der Heijden et al. 2015). Yet, to properly validate such a dramatic climate change-carbon feedback loop and to unravel the role lianas play therein, one needs to consider the interplay between the energy, the water, and the carbon cycles. Most studies that investigated the role of lianas have neglected those interactions, primarily focusing on a single biogeochemical cycle. As illustrated here, calibrated vegetation models are key to reveal the relative magnitude of all the interacting and contradictory impacts lianas have on forests, as well as the feedbacks they generate.

While lianas remain largely ignored by vegetation models, we demonstrate here that they profoundly alter the energy balance and the carbon cycle of tropical forests, which is *per se* a strong argument to take them into account when predicting the future land carbon sink.

## Acknowledgements

This research was funded by the European Research Council Starting Grant 637643 (TREECLIMBERS). The computational resources and services used in this work were provided by the VSC (Flemish Supercomputer Center), funded by the Research Foundation - Flanders (FWO) and the Flemish Government – department EWI. During the preparation of this manuscript, FM was first funded by the BAEF and the WBI as a research fellow and then by the FWO as a junior postdoc and is thankful to these organizations for their financial support. HPTDD was also a BAEF research fellow during the preparation of this manuscript and is as grateful to this organization for its support. MCD was supported by NSF ABI grant 1458021. We are grateful to the whole PEcAn group and the ED2 team for helpful discussions and support related to the functioning of BETY, PEcAn and ED-RTM. The research carried out at the Jet Propulsion Laboratory, California Institute of Technology, was under a contract with the National Aeronautics and Space Administration. ML was supported by the NASA Postdoctoral Program, administered by Universities Space Research Association under contract with NASA. GatorEye data collection and processing by AMAZ and ENB was supported by the McIntire-Stennis program of the USDA and the University of Florida.

## Author contribution

FM, MV, MD and HV designed the study. FM implemented the workflow, ran the simulations and processed the results with inputs and support from MD and AS for PEcAn and ED-RTM technical aspects. Some of the co-authors contributed with critical data to calibrate or validate the model outputs, including MV and HML (WorldView-3), JAG and ASA (raw spectral data), and ENB and AMAZ (GatorEye LiDAR). All authors contributed to the manuscript and critically revised it.

## Conflict of interest

None declared.

## Data and code availability

The code for ED2.2 and ED-RTM are available on Github at https://github.com/femeunier/ED2 in the ED and EDR folders, respectively (tag Liana_v.1). All data and scripts used in this study are also stored on github (https://github.com/femeunier/LianaAlbedo). GatorEye LiDAR data is available for download from www.gatoreye.org.

## Notes

### Competing Interest Statement

The authors have declared no competing interest.

## References

Arias, Paola A., Rong Fu, Carlos D. Hoyos, Wenhong Li, and Liming Zhou. 2011. “Changes in Cloudiness over the Amazon Rainforests during the Last Two Decades: Diagnostic and Potential Causes.” Climate Dynamics 37 (5): 1151–64. https://doi.org/10.1007/s00382-010-0903-2.

Asner, Gregory P. 2008. “Hyperspectral Remote Sensing of Canopy Chemistry, Physiology, and Biodiversity in Tropical Rainforests.” Hyperspectral Remote Sensing of Tropical and Sub-Tropical Forests, January, 261–96.

Asner, Gregory P., and Roberta E. Martin. 2012. “Contrasting Leaf Chemical Traits in Tropical Lianas and Trees: Implications for Future Forest Composition.” Ecology Letters 15 (9): 1001–7. https://doi.org/10.1111/j.1461-0248.2012.01821.x.

Avalos, Gerardo, and Stephen S. Mulkey. 1999. “Seasonal Changes in Liana Cover in the Upper Canopy of a Neotropical Dry Forest.” Biotropica 31 (1): 186–92. https://doi.org/10.2307/2663973.

Avitabile, Valerio, Martin Herold, Gerard B. M. Heuvelink, Simon L. Lewis, Oliver L. Phillips, Gregory P. Asner, John Armston, et al. 2016. “An Integrated Pan-Tropical Biomass Map Using Multiple Reference Datasets.” Global Change Biology 22 (4): 1406–20. https://doi.org/10.1111/gcb.13139.

Beer, Christian, Markus Reichstein, Enrico Tomelleri, Philippe Ciais, Martin Jung, Nuno Carvalhais, Christian Rödenbeck, et al. 2010. “Terrestrial Gross Carbon Dioxide Uptake: Global Distribution and Covariation with Climate.” Science 329 (5993): 834–38. https://doi.org/10.1126/science.1184984.

Best, M. J., M. Pryor, D. B. Clark, G. G. Rooney, R. L. H. Essery, C. B. Ménard, J. M. Edwards, et al. 2011. “The Joint UK Land Environment Simulator (JULES), Model Description – Part 1: Energy and Water Fluxes.” Geoscientific Model Development 4 (3): 677–99. https://doi.org/10.5194/gmd-4-677-2011.

Blondeel, Haben, Michael P. Perring, Leen Depauw, Emiel De Lombaerde, Dries Landuyt, Pieter De Frenne, and Kris Verheyen. 2020. “Light and Warming Drive Forest Understorey Community Development in Different Environments.” Global Change Biology 26 (3): 1681–96. https://doi.org/10.1111/gcb.14955.

Bonan, Gordon. 2019. Climate Change and Terrestrial Ecosystem Modeling. 1st ed. Cambridge University Press. https://doi.org/10.1017/9781107339217.

Bonan, Gordon B. 2008. “Forests and Climate Change: Forcings, Feedbacks, and the Climate Benefits of Forests.” Science 320 (5882): 1444–49.

Bongers, F., and F. J. (Department of Forestry Sterck. 1998. “Architecture and Development of Rainforest Trees: Responses to Light Variation.” http://agris.fao.org/agris-search/search.do?recordID=GB1997050682.

Bongers, Frans, Corneille E. N. Ewango, Masha T. van der Sande, and Lourens Poorter. 2020. “Liana Species Decline in Congo Basin Contrasts with Global Patterns.” Ecology 101 (5): e03004. https://doi.org/10.1002/ecy.3004.

Braak, Cajo J. F. ter, and Jasper A. Vrugt. 2008. “Differential Evolution Markov Chain with Snooker Updater and Fewer Chains.” Statistics and Computing 18 (4): 435–46. https://doi.org/10.1007/s11222-008-9104-9.

Brinck, Katharina, Rico Fischer, Jürgen Groeneveld, Sebastian Lehmann, Mateus Dantas De Paula, Sandro Pütz, Joseph O. Sexton, Danxia Song, and Andreas Huth. 2017. “High Resolution Analysis of Tropical Forest Fragmentation and Its Impact on the Global Carbon Cycle.” Nature Communications 8 (1): 1–6. https://doi.org/10.1038/ncomms14855.

Castro-Esau, K, G.A. Sanchez-Azofeifa, and T. Caelli. 2004. “Discrimination of Lianas and Trees with Leaf-Level Hyperspectral Data.” Remote Sensing of Environment 90 (3): 353–72. https://doi.org/10.1016/j.rse.2004.01.013.

Chen, Ya Jun, Stefan A. Schnitzer, Yong Jiang Zhang, Ze Xin Fan, Guillermo Goldstein, Kyle W. Tomlinson, Hua Lin, Jiao Lin Zhang, and Kun Fang Cao. 2017. “Physiological Regulation and Efficient Xylem Water Transport Regulate Diurnal Water and Carbon Balances of Tropical Lianas.” Functional Ecology 31 (2): 306–17. https://doi.org/10.1111/1365-2435.12724.

Chen, Ya-Jun, Kun-Fang Cao, Stefan A. Schnitzer, Ze-Xin Fan, Jiao-Lin Zhang, and Frans Bongers. 2015. “Water-Use Advantage for Lianas over Trees in Tropical Seasonal Forests.” New Phytologist 205 (1): 128–36. https://doi.org/10.1111/nph.13036.

Condit, Richard, Rolando Pérez, Salomon Aguilar, Suzanne Lao, Robin Foster, and Stephen Hubbell. 2019. “Complete Data from the Barro Colorado 50-Ha Plot: 423617 Trees, 35 Years.” Smithsonian Center for Tropical Forest Science. https://doi.org/10.15146/5xcp-0d46.

Cooley, T., G.P. Anderson, G.W. Felde, M.L. Hoke, A.J. Ratkowski, J.H. Chetwynd, J.A. Gardner, et al. 2002. “FLAASH, a MODTRAN4-Based Atmospheric Correction Algorithm, Its Application and Validation.” In IEEE International Geoscience and Remote Sensing Symposium, 3:1414–18. Toronto, Ont., Canada: IEEE. https://doi.org/10.1109/IGARSS.2002.1026134.

De Lombaerde, Emiel, Haben Blondeel, Lander Baeten, Dries Landuyt, Michael P. Perring, Leen Depauw, Sybryn L. Maes, Bin Wang, and Kris Verheyen. 2020. “Light, Temperature and Understorey Cover Predominantly Affect Early Life Stages of Tree Seedlings in a Multifactorial Mesocosm Experiment.” Forest Ecology and Management 461 (April): 117907. https://doi.org/10.1016/j.foreco.2020.117907.

Detto, Matteo, S. Joseph Wright, Osvaldo Calderón, and Helene C. Muller-Landau. 2018. “Resource Acquisition and Reproductive Strategies of Tropical Forest in Response to the El Niño–Southern Oscillation.” Nature Communications 9 (1): 1–8. https://doi.org/10.1038/s41467-018-03306-9.

Dewalt, Saara J., Stefan A. Schnitzer, Luciana F. Alves, Frans Bongers, Robyn J. Burnham, Zhiquan Cai, Walter P. Carson, et al. 2014. “Biogeographical Patterns of Liana Abundance and Diversity.” Ecology of Lianas, 131–46. https://doi.org/10.1002/9781118392409.ch11.

Dietze, Michael C., Shawn P. Serbin, Carl Davidson, Ankur R. Desai, Xiaohui Feng, Ryan Kelly, Rob Kooper, et al. 2014. “A Quantitative Assessment of a Terrestrial Biosphere Model’s Data Needs across North American Biomes.” Journal of Geophysical Research: Biogeosciences 119 (3): 286–300. https://doi.org/10.1002/2013JG002392.

Dietze, Michael C., Michael S. Wolosin, and James S. Clark. 2008. “Capturing Diversity and Interspecific Variability in Allometries: A Hierarchical Approach.” Forest Ecology and Management 256 (11): 1939–48. https://doi.org/10.1016/j.foreco.2008.07.034.

Doughty, Christopher E., Paul Efren Santos-Andrade, Alexander Shenkin, Gregory R. Goldsmith, Lisa P. Bentley, Benjamin Blonder, Sandra Díaz, et al. 2018. “Tropical Forest Leaves May Darken in Response to Climate Change.” Nature Ecology & Evolution 2 (12): 1918–24. https://doi.org/10.1038/s41559-018-0716-y.

Elzhov, Timur V., Katharine M. Mullen, Andrej-Nikolai Spiess, and Ben Bolker. 2016. Minpack.Lm: R Interface to the Levenberg-Marquardt Nonlinear Least-Squares Algorithm Found in MINPACK, Plus Support for Bounds (version 1.2-1). https://CRAN.R-project.org/package=minpack.lm.

Estrada-Villegas, Sergio, and Stefan A. Schnitzer. 2018. “A Comprehensive Synthesis of Liana Removal Experiments in Tropical Forests.” Biotropica 50 (5): 729–39. https://doi.org/10.1111/btp.12571.

Falster, Daniel S., Remko A. Duursma, Masae I. Ishihara, Diego R. Barneche, Richard G. FitzJohn, Angelica Vårhammar, Masahiro Aiba, et al. 2015. “BAAD: A Biomass And Allometry Database for Woody Plants.” Ecology 96 (5): 1445–1445. https://doi.org/10.1890/14-1889.1.

Fischlin, Andreas, G. F. Midgley, Jeff Price, Rik Leemans, Brij Gopal, Carol Turley, Mark Rounsevell, Pauline Dube, Juan Tarazona, and Andrei Velichko. 2007. “Ecosystems, Their Properties, Goods, and Services.” In: Parry, M.L., Canziani, O.F., Palutikof, J.P., van Der Linden, P.J., Hanson, C.E., Eds. Climate Change 2007: Impacts, Adaptation and Vulnerability. Contribution of Working Group II to the Fourth Assessment Report of the Intergovernmental Panel on Climate Change. Cambridge, UK: Cambridge University Press: 211-272., 211–72.

Fisher, Joshua B., Deborah N. Huntzinger, Christopher R. Schwalm, and Stephen Sitch. 2014. “Modeling the Terrestrial Biosphere.” Annual Review of Environment and Resources 39 (1): 91–123. https://doi.org/10.1146/annurev-environ-012913-093456.

Fisher, Rosie A., Charles D. Koven, William R. L. Anderegg, Bradley O. Christoffersen, Michael C. Dietze, Caroline E. Farrior, Jennifer A. Holm, et al. 2018. “Vegetation Demographics in Earth System Models: A Review of Progress and Priorities.” Global Change Biology 24 (1): 35–54. https://doi.org/10.1111/gcb.13910.

Fisher, Rosie, Nate McDowell, Drew Purves, Paul Moorcroft, Stephen Sitch, Peter Cox, Chris Huntingford, Patrick Meir, and F. Ian Woodward. 2010. “Assessing Uncertainties in a Second-Generation Dynamic Vegetation Model Caused by Ecological Scale Limitations.” New Phytologist 187 (3): 666–81. https://doi.org/10.1111/j.1469-8137.2010.03340.x.

Fu, Rong, Lei Yin, Wenhong Li, Paola A. Arias, Robert E. Dickinson, Lei Huang, Sudip Chakraborty, et al. 2013. “Increased Dry-Season Length over Southern Amazonia in Recent Decades and Its Implication for Future Climate Projection.” Proceedings of the National Academy of Sciences 110 (45): 18110–15. https://doi.org/10.1073/pnas.1302584110.

Gehring, C, S Park, and M Denich. 2004. “Liana Allometric Biomass Equations for Amazonian Primary and Secondary Forest.” Forest Ecology and Management 195 (1–2): 69–83. https://doi.org/10.1016/j.foreco.2004.02.054.

Gerwing, Jeffrey John, and Damião Lopes Farias. 2000. “Integrating Liana Abundance and Forest Stature into an Estimate of Total Aboveground Biomass for an Eastern Amazonian Forest.” Journal of Tropical Ecology 16 (3): 327–35. https://doi.org/10.1017/S0266467400001437.

Goudriaan, J. 1977. “Crop Micrometeorology : A Simulation Study.” Phd, Wageningen: Pudoc. https://library.wur.nl/WebQuery/wurpubs/70980.

Guzmán Q., J. Antonio, Benoit Rivard, and G. Arturo Sánchez-Azofeifa. 2018. “Discrimination of Liana and Tree Leaves from a Neotropical Dry Forest Using Visible-near Infrared and Longwave Infrared Reflectance Spectra.” Remote Sensing of Environment 219 (December): 135–44. https://doi.org/10.1016/j.rse.2018.10.014.

Hartig, Florian, Francesco Minunno, Stefan Paul, David Cameron, Tankred Ott, and Maximilian Pichler. 2019. BayesianTools: General-Purpose MCMC and SMC Samplers and Tools for Bayesian Statistics (version 0.1.7). https://CRAN.R-project.org/package=BayesianTools.

Heijden, Geertje M. van der, Stefan a. Schnitzer, Jennifer S. Powers, and Oliver L. Phillips. 2013. “Liana Impacts on Carbon Cycling, Storage and Sequestration in Tropical Forests.” Biotropica 45 (6): 682–92. https://doi.org/10.1111/btp.12060.

Heijden, Geertje van der, Jennifer S Powers, and S.A. Schnitzer. 2015. “Lianas Reduce Forest-Level Carbon Accumulation and Storage.” PNAS 112 (43): 13267–71. https://doi.org/10.1073/pnas.1504869112.

Jacquemoud, S, C Bacour, H Poilvé, and J. -P Frangi. 2000. “Comparison of Four Radiative Transfer Models to Simulate Plant Canopies Reflectance: Direct and Inverse Mode.” Remote Sensing of Environment 74 (3): 471–81. https://doi.org/10.1016/S0034-4257(00)00139-5.

Kalacska, M., S. Bohlman, G. A. Sanchez-Azofeifa, K. Castro-Esau, and T. Caelli. 2007. “Hyperspectral Discrimination of Tropical Dry Forest Lianas and Trees: Comparative Data Reduction Approaches at the Leaf and Canopy Levels.” Remote Sensing of Environment 109 (4): 406–15. https://doi.org/10.1016/j.rse.2007.01.012.

Kazda, M, and J Salzer. 2000. “Leaves of Lianas and Self-Supporting Plants Differ in Mass per Unit Area and in Nitrogen Content.” Plant Biology 2: 268–71.

Konapala, Goutam, Ashok K. Mishra, Yoshihide Wada, and Michael E. Mann. 2020. “Climate Change Will Affect Global Water Availability through Compounding Changes in Seasonal Precipitation and Evaporation.” Nature Communications 11 (1): 3044. https://doi.org/10.1038/s41467-020-16757-w.

Koricheva, Julia, Jessica Gurevitch, and Kerrie Mengersen, eds. 2013. Handbook of Meta-Analysis in Ecology and Evolution. 2013th edition. Princeton: Princeton University Press.

Krishna Moorthy, Sruthi M. 2019. “Assessing the Role of Lianas in Tropical Forest Structure with Terrestrial LiDAR.” Dissertation, Ghent University. http://hdl.handle.net/1854/LU-8636809.

Kurzel, Brian P., Stefan a. Schnitzer, and Walter P. Carson. 2006. “Predicting Liana Crown Location from Stem Diameter in Three Panamanian Lowland Forests.” Biotropica 38 (2): 262–66. https://doi.org/10.1111/j.1744-7429.2006.00135.x.

Lawrence, David M., Rosie A. Fisher, Charles D. Koven, Keith W. Oleson, Sean C. Swenson, Gordon Bonan, Nathan Collier, et al. 2019. “The Community Land Model Version 5: Description of New Features, Benchmarking, and Impact of Forcing Uncertainty.” Journal of Advances in Modeling Earth Systems 11 (12): 4245–87. https://doi.org/10.1029/2018MS001583.

LeBauer, David S., Dan Wang, Katherine T. Richter, Carl C. Davidson, and Michael C. Dietze. 2013. “Facilitating Feedbacks between Field Measurements and Ecosystem Models.” Ecological Monographs 83 (2): 133–154.

Liou, K.N. 2002. An Introduction to Atmospheric Radiation. Vol. 84. International Geophysics Series. https://books.google.com/books/about/An_Introduction_to_Atmospheric_Radiation.html?id=mQ1DiDpX34UC.

Longo, Marcos, Ryan G. Knox, Naomi M. Levine, Abigail L. S. Swann, David M. Medvigy, Michael C. Dietze, Yeonjoo Kim, et al. 2019a. “The Biophysics, Ecology, and Biogeochemistry of Functionally Diverse, Vertically-and Horizontally-Heterogeneous Ecosystems: The Ecosystem Demography Model, Version 2.2 – Part 2: Model Evaluation.” Geoscientific Model Development Discussions, March, 1–34. https://doi.org/10.5194/gmd-2019-71.

Longo, Marcos, Ryan G. Knox, Naomi M. Levine, Abigail L. S. Swann, David M. Medvigy, Michael C. Dietze, Yeonjoo Kim. 2019b. “The Biophysics, Ecology, and Biogeochemistry of Functionally Diverse, Vertically and Horizontally Heterogeneous Ecosystems: The Ecosystem Demography Model, Version 2.2 – Part 2: Model Evaluation for Tropical South America.” Geoscientific Model Development 12 (10): 4347–74. https://doi.org/10.5194/gmd-12-4347-2019.

Longo, Marcos, Sassan Saatchi, Michael Keller, Kevin Bowman, António Ferraz, Paul R. Moorcroft, Douglas C. Morton, et al. 2020. “Impacts of Degradation on Water, Energy, and Carbon Cycling of the Amazon Tropical Forests.” Journal of Geophysical Research: Biogeosciences 125 (8): e2020JG005677. https://doi.org/10.1029/2020JG005677.

Marvin, David C., Gregory P. Asner, and Stefan A. Schnitzer. 2016. “Liana Canopy Cover Mapped throughout a Tropical Forest with High-Fidelity Imaging Spectroscopy.” Remote Sensing of Environment 176: 98–106. https://doi.org/10.1016/j.rse.2015.12.028.

Medvigy, D., S. C. Wofsy, J. W. Munger, D. Y. Hollinger, and P. R. Moorcroft. 2009. “Mechanistic Scaling of Ecosystem Function and Dynamics in Space and Time: Ecosystem Demography Model Version 2.” Journal of Geophysical Research 114 (G1). https://doi.org/10.1029/2008JG000812.

Meroni, M., R. Colombo, and C. Panigada. 2004. “Inversion of a Radiative Transfer Model with Hyperspectral Observations for LAI Mapping in Poplar Plantations.” Remote Sensing of Environment 92 (2): 195–206. https://doi.org/10.1016/j.rse.2004.06.005.

Meunier, Félicien, Hans Verbeeck, Betsy Cowdery, Stefan A. Schnitzer, Chris M. Smith-Martin, Jennifer Powers, Xiangtao Xu, et al. 2020. “Unraveling the Relative Role of Light and Water Competition between Lianas and Trees in Tropical Forests.” Journal of Ecology, October, 1365-2745.13540. https://doi.org/10.1111/1365-2745.13540.

Mitchard, Edward T. A. 2018. “The Tropical Forest Carbon Cycle and Climate Change.” Nature 559 (7715): 527–34. https://doi.org/10.1038/s41586-018-0300-2.

Oleson, Keith, M. Lawrence, B. Bonan, Beth Drewniak, Maoyi Huang, D. Koven, Samuel Levis, et al. 2013. “Technical Description of Version 4.5 of the Community Land Model (CLM).” https://doi.org/10.5065/D6RR1W7M.

Pan, Y., R. A. Birdsey, J. Fang, R. Houghton, P. E. Kauppi, W. A. Kurz, O. L. Phillips, et al. 2011. “A Large and Persistent Carbon Sink in the World’s Forests.” Science 333 (6045): 988–93. https://doi.org/10.1126/science.1201609.

Park, John Y., Helene C. Muller-Landau, Jeremy W. Lichstein, Sami W. Rifai, Jonathan P. Dandois, and Stephanie A. Bohlman. 2019. “Quantifying Leaf Phenology of Individual Trees and Species in a Tropical Forest Using Unmanned Aerial Vehicle (UAV) Images.” Remote Sensing 11 (13): 1534. https://doi.org/10.3390/rs11131534.

Pau, Stephanie, Matteo Detto, Youngil Kim, and Christopher J. Still. 2018. “Tropical Forest Temperature Thresholds for Gross Primary Productivity.” Ecosphere 9 (7): e02311. https://doi.org/10.1002/ecs2.2311.

Peng Gong, Ruiliang Pu, G.S. Biging, and M.R. Larrieu. 2003. “Estimation of Forest Leaf Area Index Using Vegetation Indices Derived from Hyperion Hyperspectral Data.” IEEE Transactions on Geoscience and Remote Sensing 41 (6): 1355–62. https://doi.org/10.1109/TGRS.2003.812910.

Phillips, O. L. 2002. “Increasing Dominance of Large Lianas in Amazonian Forests.” Nature 418 (6899): 770–74. https://doi.org/10.1038/nature00926.

Piao, Shilong, Xuhui Wang, Taejin Park, Chi Chen, Xu Lian, Yue He, Jarle W. Bjerke, et al. 2020. “Characteristics, Drivers and Feedbacks of Global Greening.” Nature Reviews Earth & Environment 1 (1): 14–27. https://doi.org/10.1038/s43017-019-0001-x.

Poorter, Lourens, Frans Bongers, Frank J. Sterck, and Hannsjörg Wöll. 2003. “Architecture of 53 Rain Forest Tree Species Differing in Adult Stature and Shade Tolerance.” Ecology 84 (3): 602–8. https://doi.org/10.1890/0012-9658(2003)084[0602:AORFTS]2.0.CO;2.

Porcia e Brugnera, Manfredo di, Félicien Meunier, Marcos Longo, Sruthi M. Krishna Moorthy, Hannes De Deurwaerder, Stefan A. Schnitzer, Damien Bonal, Boris Faybishenko, and Hans Verbeeck. 2019. “Modeling the Impact of Liana Infestation on the Demography and Carbon Cycle of Tropical Forests.” Global Change Biology 25 (11): 3767–80. https://doi.org/10.1111/gcb.14769.

Powell, Thomas, Lara Kueppers, and Steve Paton. 2017. “Seven Years (2008-2014) of Meteorological Observations plus a Synthetic El Nino Drought for BCI Panama.” Next-Generation Ecosystem Experiments Tropics; Lawrence Berkeley National Laboratory (See acknowledgements below for the source of the original raw data). https://doi.org/10.15486/ngt/1414275.

Powell, Thomas L., David R. Galbraith, Bradley O. Christoffersen, Anna Harper, Hewlley M. A. Imbuzeiro, Lucy Rowland, Samuel Almeida, et al. 2013. “Confronting Model Predictions of Carbon Fluxes with Measurements of Amazon Forests Subjected to Experimental Drought.” The New Phytologist 200 (2): 350–65. https://doi.org/10.1111/nph.12390.

Powell, Thomas L., Charles D. Koven, Daniel J. Johnson, Boris Faybishenko, Rosie A. Fisher, Ryan G. Knox, Nate G. McDowell, et al. 2018. “Variation in Hydroclimate Sustains Tropical Forest Biomass and Promotes Functional Diversity.” New Phytologist 219 (3): 932–46. https://doi.org/10.1111/nph.15271.

Purves, Drew, and Stephen Pacala. 2008. “Predictive Models of Forest Dynamics.” Science 320 (5882): 1452–53. https://doi.org/10.1126/science.1155359.

Putz, Francis E. 1983. “Liana Biomass and Leaf Area of a “Tierra Firme “Forest in the Rio Negro Basin, Venezuela.” Biotropica 15 (3): 185–89.

Reis, Simone Matias, Beatriz Schwantes Marimon, Paulo S. Morandi, Fernando Elias, Adriane Esquivel-Muelbert, Ben Hur Marimon Junior, Sophie Fauset, et al. 2020. “Causes and Consequences of Liana Infestation in Southern Amazonia.” Journal of Ecology 108 (6): 2184–97. https://doi.org/10.1111/1365-2745.13470.

Sánchez-Azofeifa, G. A., and K. Castro-Esau. 2006. “Canopy Observations on the Hyperspectral Properties of a Community of Tropical Dry Forest Lianas and Their Host Trees.” International Journal of Remote Sensing 27 (10): 2101–9. https://doi.org/10.1080/01431160500444749.

Sánchez-Azofeifa, G. Arturo, Karen Castro, S. Joseph Wright, John Gamon, Margaret Kalacska, Benoit Rivard, Stefan A. Schnitzer, and Ji L. Feng. 2009. “Differences in Leaf Traits, Leaf Internal Structure, and Spectral Reflectance between Two Communities of Lianas and Trees: Implications for Remote Sensing in Tropical Environments.” Remote Sensing of Environment 113 (10): 2076–88. https://doi.org/10.1016/j.rse.2009.05.013.

Sande, Masha T. van der, Lourens Poorter, Stefan A. Schnitzer, and Lars Markesteijn. 2013. “Are Lianas More Drought-Tolerant than Trees? A Test for the Role of Hydraulic Architecture and Other Stem and Leaf Traits.” Oecologia 172 (4): 961–72. https://doi.org/10.1007/s00442-012-2563-x.

Schnitzer, Stefan A. 2005. “A Mechanistic Explanation for Global Patterns of Liana Abundance and Distribution.” The American Naturalist 166 (2): 262–76. https://doi.org/10.1086/431250.

Schnitzer, Stefan A., and Frans Bongers. 2002. “The Ecology of Lianas and Their Role in Forests.” Trends in Ecology & Evolution 17 (5): 223–30. https://doi.org/10.1016/S0169-5347(02)02491-6.

Schnitzer, Stefan A., and Frans Bongers. 2011. “Increasing Liana Abundance and Biomass in Tropical Forests: Emerging Patterns and Putative Mechanisms.” Ecology Letters 14 (4): 397–406. https://doi.org/10.1111/j.1461-0248.2011.01590.x.

Schnitzer, Stefan A., Saara J. DeWalt, and Jerome Chave. 2006. “Censusing and Measuring Lianas: A Quantitative Comparison of the Common Methods1.” Biotropica 38 (5): 581–91. https://doi.org/10.1111/j.1744-7429.2006.00187.x.

Schnitzer, Stefan A., and Geertje M. F. van der Heijden. 2019. “Lianas Have a Seasonal Growth Advantage over Co-occurring Trees.” Ecology 100 (5): e02655. https://doi.org/10.1002/ecy.2655.

Schnitzer, Stefan A., Mirjam E. Kuzee, and Frans Bongers. 2005. “Disentangling Above-and below-Ground Competition between Lianas and Trees in a Tropical Forest.” Journal of Ecology 93 (6): 1115–25. https://doi.org/10.1111/j.1365-2745.2005.01056.x.

Schnitzer, Stefan a, Scott a Mangan, James W Dalling, Claire a Baldeck, Stephen P Hubbell, Alicia Ledo, Helene Muller-Landau, et al. 2012. “Liana Abundance, Diversity, and Distribution on Barro Colorado Island, Panama.” PloS One 7 (12): e52114–e52114. https://doi.org/10.1371/journal.pone.0052114.

Selaya, N. G., and N. P R Anten. 2008. “Differences in Biomass Allocation, Light Interception and Mechanical Stability between Lianas and Trees in Early Secondary Tropical Forest.” Functional Ecology 22: 30–39. https://doi.org/10.1111/j.1365-2435.2007.01350.x.

Sellers, P. J. 1985. “Canopy Reflectance, Photosynthesis and Transpiration.” International Journal of Remote Sensing 6 (8): 1335–72. https://doi.org/10.1080/01431168508948283.

Serbin, Shawn P., Aditya Singh, Ankur R. Desai, Sean G. Dubois, Andrew D. Jablonski, Clayton C. Kingdon, Eric L. Kruger, and Philip A. Townsend. 2015. “Remotely Estimating Photosynthetic Capacity, and Its Response to Temperature, in Vegetation Canopies Using Imaging Spectroscopy.” Remote Sensing of Environment 167 (September): 78–87. https://doi.org/10.1016/j.rse.2015.05.024.

Shiklomanov, Alexey N., Ben Bond-Lamberty, Jeff W. Atkins, and Christopher M. Gough. 2020. “Structure and Parameter Uncertainty in Centennial Projections of Forest Community Structure and Carbon Cycling.” Global Change Biology, August, gcb.15164. https://doi.org/10.1111/gcb.15164.

Shiklomanov, Alexey N., Michael C. Dietze, Istem Fer, Toni Viskari, and Shawn P. Serbin. 2020. “Cutting out the Middleman: Calibrating and Validating a Dynamic Vegetation Model (ED2-PROSPECT5) Using Remotely Sensed Surface Reflectance.” Geoscientific Model Development Discussions, October, 1–35. https://doi.org/10.5194/gmd-2020-324.

Shiklomanov, Alexey N., Michael C. Dietze, Toni Viskari, Philip A. Townsend, and Shawn P. Serbin. 2016. “Quantifying the Influences of Spectral Resolution on Uncertainty in Leaf Trait Estimates through a Bayesian Approach to RTM Inversion.” Remote Sensing of Environment 183 (September): 226–38. https://doi.org/10.1016/j.rse.2016.05.023.

Smith-Martin, Chris M., Xiangtao Xu, David Medvigy, Stefan A. Schnitzer, and Jennifer S. Powers. 2019. “Allometric Scaling Laws Linking Biomass and Rooting Depth Vary across Ontogeny and Functional Groups in Tropical Dry Forest Lianas and Trees.” New Phytologist n/a (n/a). https://doi.org/10.1111/nph.16275.

Spracklen, D.V., J.C.A. Baker, L. Garcia-Carreras, and J.H. Marsham. 2018. “The Effects of Tropical Vegetation on Rainfall.” Annual Review of Environment and Resources 43 (1): 193–218. https://doi.org/10.1146/annurev-environ-102017-030136.

Stevens, George C. 1987. “Lianas as Structural Parasites : The Bursera Simaruba Example Published by : Ecological Society of America Linked References Are Available on JSTOR for This Article :” Ecology 68 (1): 77–81.

Verbeeck, Hans, and Elizabeth Kearsley. 2016. “The Importance of Including Lianas in Global Vegetation Models.” Proceedings of the National Academy of Sciences of the United States of America 113 (1): E4. https://doi.org/10.1073/pnas.1521343113.

Viskari, Toni, Alexey Shiklomanov, Michael C. Dietze, and Shawn P. Serbin. 2019. “The Influence of Canopy Radiation Parameter Uncertainty on Model Projections of Terrestrial Carbon and Energy Cycling.” PLOS ONE 14 (7): e0216512. https://doi.org/10.1371/journal.pone.0216512.

Xu, Xiangtao, David Medvigy, Jennifer S. Powers, Justin M. Becknell, and Kaiyu Guan. 2016. “Diversity in Plant Hydraulic Traits Explains Seasonal and Inter-Annual Variations of Vegetation Dynamics in Seasonally Dry Tropical Forests.” New Phytologist 212 (1): 80–95. https://doi.org/10.1111/nph.14009.

Yuan, Chun ming, Tao Wu, Yun fen Geng, Yong Chai, and Jia bo Hao. 2016. “Phenotypic Plasticity of Lianas in Response to Altered Light Environment.” Ecological Research 31 (3): 375–84. https://doi.org/10.1007/s11284-016-1343-1.

Yuan, Xu, Kati Laakso, Philip Marzahn, and G. Arturo Sanchez-Azofeifa. 2019. “Canopy Temperature Differences between Liana-Infested and Non-Liana Infested Areas in a Neotropical Dry Forest.” Forests 10 (10): 890. https://doi.org/10.3390/f10100890.

