## Supplementary material for "Liana optical traits increase tropical forest albedo and reduce ecosystem productivity"

#### Appendix A: The radiative transfer model of ED2

In this appendix, we describe how the parameters of Table 1 (main text) translate in Equations 45-47 from Longo et al. (2019a), who describe in detail the radiation model of ED2.2. Each cohort *k* is assumed to be one single layer of vegetation of effective plant area index $\Phi_k$ with constant and homogeneous optical properties. The effective plant area index of cohort *k* is defined as the sum of the wood area and the effective leaf areas:

$$\Phi_k = \Omega \cdot LAI_k + WAI_k$$

where $LAI_k$ and $WAI_k$ are the leaf and wood area index of cohort *k* and Ω is the PFT-specific clumping factor.

At any time, the top boundary condition of the radiative transfer model is provided by the current forcing data. The visible and solar infrared downward irradiance is composed of a direct (beam) and diffuse (isotropic) components. When intercepted by the cohorts, direct irradiance can be either backscattered or forward-scattered as diffuse radiation. Following Sellers (1985), the extinction of downward direct irradiance (Eq. A1) and the two-stream model for hemispheric diffuse irradiance (Eqs. A2-A3) for each wavelength is given by :

$$\underbrace{\mu^{\odot}_k \frac{d\dot{Q}^{\odot}_{\lambda k}}{d\Phi}}_{Downward\ direct\ profile} = \underbrace{-\dot{Q}^{\odot}_{\lambda k}}_{Interception} \qquad \text{Eq. A1}$$

$$\underbrace{\overline{\mu}_k \frac{d\dot{Q}^{\Downarrow}_{\lambda k}}{d\Phi}}_{Downward\ diffuse\ profile} = \underbrace{-\dot{Q}^{\Downarrow}_{\lambda k}}_{Interception} + \underbrace{\left(1-\beta_{\lambda k}\right)\cdot\varsigma_{\lambda k}\cdot\dot{Q}^{\Downarrow}_{\lambda k}}_{\substack{Forward\ scattering \\ (downward\ diffuse)}} + \underbrace{\beta_{\lambda k}\cdot\varsigma_{\lambda k}\cdot\dot{Q}^{\Uparrow}_{\lambda k}}_{\substack{Backscattering \\ (upward\ diffuse)}} + \underbrace{\frac{\overline{\mu}_k}{\mu^{\odot}_k}\left(1-\beta^{\odot}_{\lambda k}\right)\cdot\varsigma_{\lambda k}\cdot\dot{Q}}_{\substack{Forward\ scattering \\ (downward\ direct)}} \qquad \text{Eq. A2}$$

$$\underbrace{-\overline{\mu}_k \frac{d\dot{Q}^{\Uparrow}_{\lambda k}}{d\Phi}}_{Upward\ diffuse\ profile} = \underbrace{-\dot{Q}^{\Uparrow}_{\lambda k}}_{Interception} + \underbrace{\left(1-\beta_{\lambda k}\right)\cdot\varsigma_{\lambda k}\cdot\dot{Q}^{\Uparrow}_{\lambda k}}_{\substack{Forward\ scattering \\ (upward\ diffuse)}} + \underbrace{\beta_{\lambda k}\cdot\varsigma_{\lambda k}\cdot\dot{Q}^{\Downarrow}_{\lambda k}}_{\substack{Backscattering \\ (downward\ diffuse)}} + \underbrace{\frac{\overline{\mu}_k}{\mu^{\odot}_k}\beta^{\odot}_{\lambda k}\cdot\varsigma_{\lambda k}\cdot\dot{Q}^{\odot}_{\lambda k}}_{\substack{Backscattering \\ (downward\ direct)}} \qquad \text{Eq. A3}$$

As compared to equations 45-47 from Longo et al. (2019), the emission terms have vanished as we only focus on the short wavelengths and emissions are negligible in these spectral domains. In addition, we replaced the wide bands *m* in the original publication by specific wavelengths λ. In Equations A1-A3, $\varsigma$ and $\beta$ are the scattering and backscattering coefficients, $\odot$ is the symbol for direct radiation, $\Downarrow$ and $\Uparrow$ the symbols for downward and upward diffuse radiations respectively, $\dot{Q}$ is irradiance, $\mu$ refers to the inverse of the optical depth, and $\Phi$ is the effective cumulative plant area index, assumed zero at the very top, and increasing downwards. $\bar{\mu}_k$ and $\mu_k^{\odot}$ are the inverse of the optical depth per unit of effective plant area index for direct and diffuse radiation, respectively and $(1 - \beta_{\lambda k})$ is the absorptivity.

For a radiation beam coming from a given angle of incidence *Z*, the inverse of the optical depth per unit of plant area is given according to Sellers (1985), and Oleson et al. (2013) by:

$$\mu(Z) = \frac{\cos(Z)}{E(Z)}$$

where *E(Z)* is the average projection of all leaves and branches onto the horizontal plan, further given after Goudriaan (1977) by:

$$E(Z) = \phi_1 + \phi_2 \cos(Z)$$

with $\phi_1$ and $\phi_2$ computed as:

$$\phi_1 = 0.5 - \omega(0.633 + 0.33 \cdot \omega)$$

$$\phi_2 = 0.877\left(1 - 2\phi_1\right)$$

where$\omega$is the leaf orientation parameter. For the diffuse radiation, we need to calculate the contribution of all angles and therefore the inverse of optical depth is calculated as:

$$\bar{\mu} = \int_0^{\pi/2} \frac{\cos(Z)}{E(Z)} \sin(Z)\, dZ = \frac{1}{\phi_2}\left(1 + \frac{\phi_1}{\phi_2} \ln\left(\frac{\phi_1}{\phi_1 + \phi_2}\right)\right)$$

The scattering parameters for each wavelength and cohort *k* are calculated just like in the Community Land Model (Oleson et al. 2013; Lawrence et al. 2019), which is derived from Goudriaan (1977) and Sellers (1985)

$$\varsigma_{\lambda k} = \rho_{\lambda k} + \tau_{\lambda k}$$

$$\beta_{\lambda k} = \frac{1}{2\varsigma_{\lambda k}}\left(\varsigma_{\lambda k} + \frac{1}{4}\left(\rho_{\lambda k} - \tau_{\lambda k}\right) \cdot (1 + \omega)^2\right)$$

where $\rho_{\lambda k}$ and $\tau_{\lambda k}$ are the reflectivity and transmissivity values at the wavelength λ that in our study are computed from PROSPECT-5 simulations. The direct scattering and backscattering coefficients are the same as Sellers (1985) and Oleson et al. (2013):

$$\beta_{\lambda k}^{\odot} = \frac{\bar{\mu}_k + \mu_k^{\odot}}{\bar{\mu}_k} \frac{\varsigma_{\lambda k}^{\odot}}{\varsigma_{\lambda k}}$$

$$\frac{\varsigma_{\lambda k}^{\odot}}{\varsigma_{\lambda k}} = \frac{1}{2\left(1 + \phi_2 \mu_k^{\odot}\right)} \left(1 - \frac{\phi_1 \mu_k^{\odot}}{1 + \phi_2 \mu_k^{\odot}} \ln\left(\frac{1 + \left(\phi_1 + \phi_2\right) \mu_k^{\odot}}{\phi_1 \mu_k^{\odot}}\right)\right)$$

#### Appendix B: Parameter data assimilation

The data assimilation approach we used in this study is adapted from Shiklomanov et al. (2016; 2020). We took a similar Bayesian approach to discriminate liana and tree leaf spectra using the leaf radiative model PROSPECT-5 and low and high liana coverage canopy spectra using ED-RTM.

At the leaf level, we calibrated the set of parameters $\theta = \left[N_{layers}, Cab, Car, Cw, Cm\right]$ given a set of observed reflectance values *X* according to the following mathematical scheme:

$$P(\theta, \sigma \mid X) = P(X \mid \theta, \sigma) \cdot P(\theta) \cdot P(\sigma)$$

$$P(X \mid \theta, \sigma) = N(PROSPECT5(\theta) \mid X, \sigma)$$

Eq. B1

where $P(\theta, \sigma \mid X)$ *is t*he joint probability posterior distribution of the parameter set, $PROSPECT5(\theta)$ is the modeled reflectance for a specific set of parameters θand$P(X \mid \theta, \sigma)$ is the likelihood. The residual error was assumed to be normally distributed with a mean of 0 and standard deviation σ. $P(\theta)$, and $P(\sigma)$ are parameters and standard deviation prior distributions, respectively.

At the stand level, we used a similar calibration scheme, simply updating the model used (ED-RTM) and the set of calibrated parameters:

$$\theta = \left[\theta_{soil}, \left(N_{layers}, Cab, Car, Cw, Cm, b1Bl, b2Bl, \Omega, \omega\right)_{i = 1 \cdots n_{PFT}}\right]$$

Eq. B2

where $\theta_{soil}$ is the soil water saturation used to estimate the soil reflectance spectrum and the remaining parameters are PFT-dependent (they vary for each of the $n_{PFT}$ considered PFTs). In ED-RTM, PROSPECT-5 is first run for each PFT to generate the leaf spectra which are then plugged into the canopy radiative transfer model.

The model goodness of fit after Bayesian calibration was estimated by computing several error statistical metrics, namely the root mean square error (RMSE), the bias (Bias), and the bias-corrected RMSE (SEPC), over three different regions of the spectrum: the visible (400-700 nm), the near infrared (700-1400 nm) and the shortwave infrared (1500-2500) following Shiklomanov et al. (2016):

$$RMSE = \sqrt{\frac{\sum_{i=1}^{n} \sum_{\lambda=1}^{m_i} \left(x_{i,\lambda} - \hat{x}_{i,\lambda}\right)^2}{N}}$$

Eq. B3

$$Bias = \frac{\sum_{i=1}^{n} \sum_{\lambda = 1}^{m_i} \left( x_{i,\lambda} - \hat{x}_{i,\lambda} \right)}{N}$$

$$SEPC = \sqrt{\frac{\sum_{i = 1}^{n} \sum_{\lambda = 1}^{m_i} \left( x_{i,\lambda} - \hat{x}_{i,\lambda} - Bias \right)^2}{N}}$$

where $n$ refers to the number of available studies, $m_i$ to the number of observations for the study $i$ in the considered region of the spectrum, and $N$ to the resulting total number of observations for each region of the spectrum. In Equation B3, $x$ and $\hat{x}$ are the simulated and observed reflectance values, respectively. We repeated the calculation of the statistical metrics described above considering the different PFTs (at the leaf-level) or the different levels of infestation (at the canopy level) as $x$ and $\hat{x}$ for both the simulated and the observed spectra to estimate whether the posterior distributions reproduced the actual differences and biases between scenarios.

### Appendix C: Model uncertainty analyses

PEcAn uncertainty analysis combines the variance of posterior distributions resulting from the model calibration with a model univariate sensitivity analysis. Doing so, the relative contribution of each parameter to the total predictive uncertainty can be estimated (LeBauer et al. 2013). More precisely, the model output (*e.g.,* simulated leaf and canopy reflectances in the different bands or ecosystem productivity) response to one-at-the time change of each parameter (median, median ± 1, 2, 3 standard deviations) is fitted using a Hermite cubic spline function, which allows translating model input variability into output variances. The relative parameter contribution to the overall variance is then simply the fraction of the total output variance explained by each parameter. For the uncertainty analysis of the leaf (PROSPECT-5) and canopy (ED-RTM) reflectances, the shortwave spectrum was divided into three regions, hereafter referred to as visible (400-700 nm), near infrared (700-1400 nm) and short-wave infrared (1500-2500 nm). For the uncertainty analysis of the vegetation model (ED2), we considered the calibrated model parameters (ω, Ω, b1Bl, b2Bl, $\tau_{vis}$, $\rho_{vis}$, $\tau_{IR}$, and $\rho_{IR}$) and extended the investigated output variables to the forest carbon (*e.g.,* GPP) and energy (*e.g.* understorey visible light) cycles.

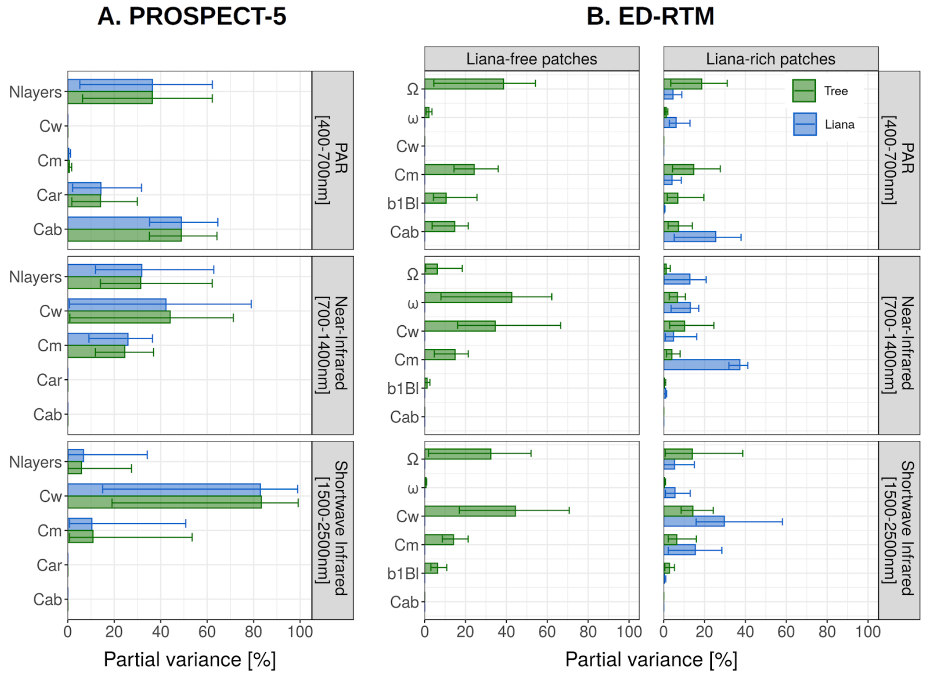


**Figure C1: Uncertainty analysis of the PROSPECT-5 (A), and ED-RTM (B) models. The bars represent the mean contributions to the overall reflectance variance generated by the different studies used to ingest leaf, canopy spectral data or both. The error bars represent the extremum contributions among studies. The uncertainty analysis was achieved on liana and tree leaves (A), or over liana-free and liana-infested patches (B). The uncertainty analyses are based on the posterior distributions (after calibration) and are shown for several output variables: visible (400-700 nm, top row), near-infrared (700-1400 nm, middle row) and short-wave infrared (1500-2500 nm, bottom row) reflectance.**

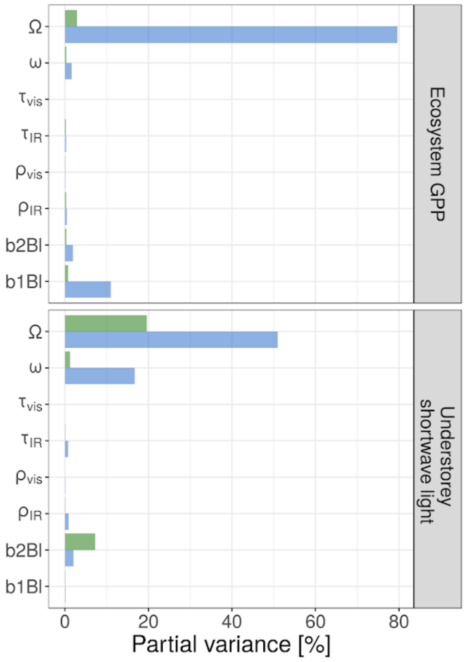


**Figure C2: Uncertainty analysis of the ED2.2 model. The uncertainty analysis was achieved over the five years of the vegetation model simulations based on the aggregated posterior distributions (after calibration). Results are shown for the ecosystem GPP (top), and the light available in the understorey (bottom).**

With regards to leaf reflectance, liana and tree leaves were both sensitive to the number of mesophyll layers (mean partial variances of 36% for lianas and trees in the visible; respectively 32% and 31% in the near-infrared, and 7% and 6% in the shortwave infrared), see Figure C1. In addition, in each region of the spectrum and for both PFTs, $N_{layers}$ was complemented by mainly two uncertain parameters: *Cab* and *Car* in the visible (49% and 14%, respectively when averaging out lianas and trees), $C_w$ and $C_m$ in the near-infrared (43% and 25%) and in the shortwave infrared (83% and 11%).

At the canopy level, while liana parameters contribute little to variance decomposition in patches with low levels of liana coverage, they became much more critical parameters in liana-rich patches, summing up on average 45%, 61% and 77% of the variance in the PAR, NIR and SWIR respectively (Figure C1). Similarly to the leaf-level, PROSPECT-5 parameters played an important role in the uncertainty according to the region of interest (*Cab* in the visible, $C_w$ and $C_m$ in the infrared), with a generally stronger impact of the leaf dry mass $C_m$ as it also determined plant *LAI*. In addition, canopy clumping (22% across all outputs and studies) and leaf orientation (13%) substantially contributed to the overall output uncertainty, with peaks of importance in the PAR and SWIR for $\Omega$ (31% and 26%, respectively, across scenarios) and in the NIR for $\omega$ (31%). These results indicated that ED-RTM calibrations were sensitive to those canopy structural traits.

In ED2, the clumping factor, especially the one of the liana PFT, drove the overall model uncertainty of the forest carbon and energy cycles, respectively illustrated by the ecosystem GPP and understorey light in Figure C2. This was explained by the fact that clumping factor posterior distribution was wide (coefficient of variation of 20%) and the model was very sensitive to its change (model elasticity of 26%). Other sensitive parameters were either more constrained (*b1Bl* and *b2Bl*), less critical for investigated model outputs ($\omega$, $\rho_{vis}$, $\tau_{vis}$) or both ($\rho_{IR}$, $\tau_{IR}$).

### Appendix D: Meta-analysis data

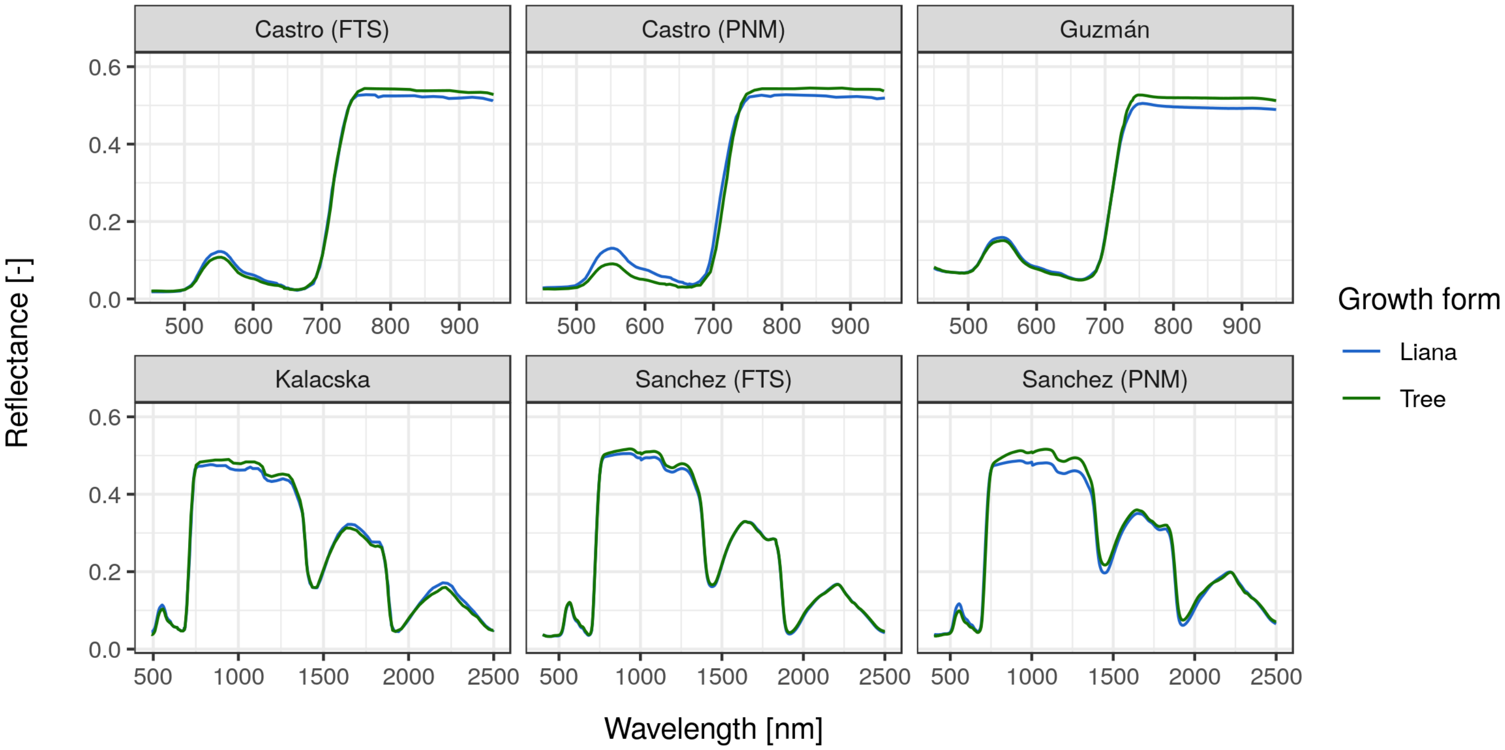


**Figure D1: Mean leaf reflectance spectra of lianas (blue) and trees (green) as observed in the different studies/sites collected through the meta-analysis.**

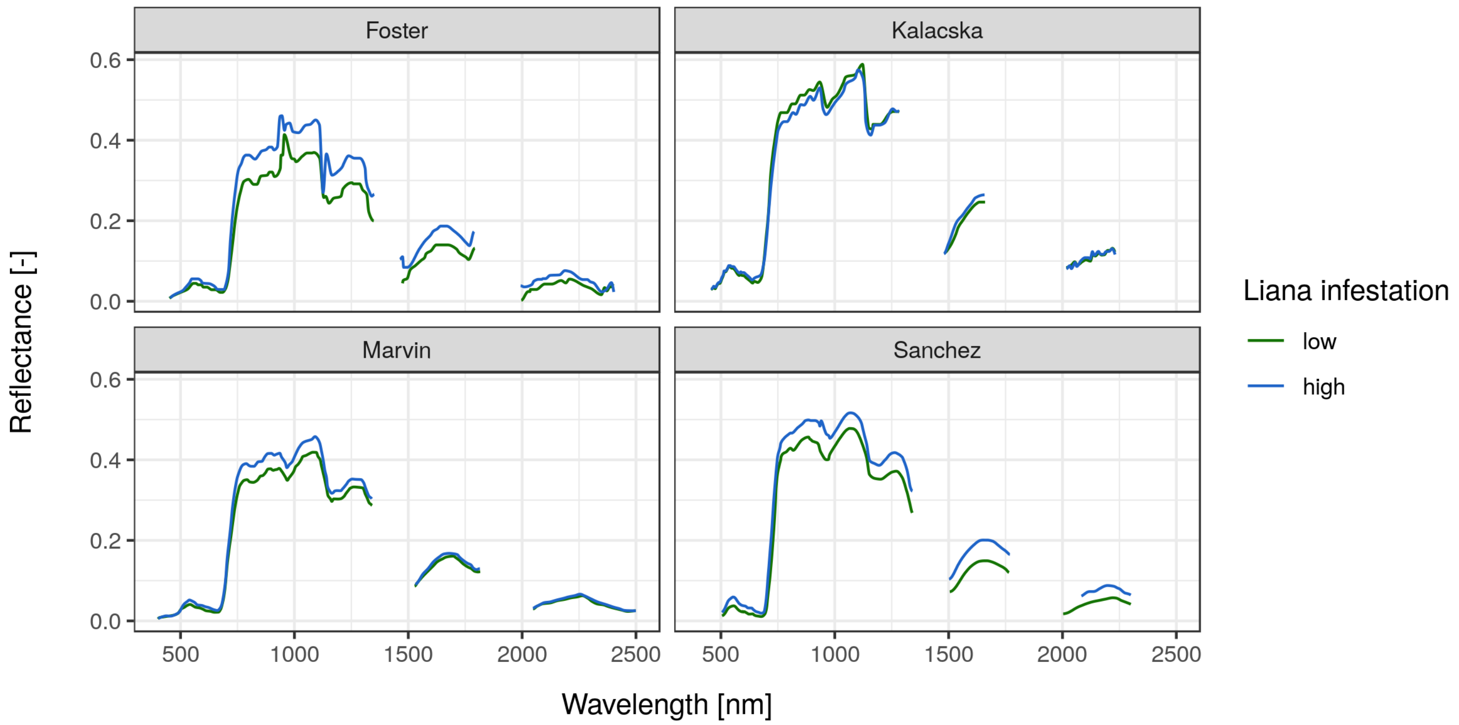


**Figure D2: Mean canopy reflectance spectra of liana-infested (blue) and liana-free (green) forest stands as observed in the different studies collected in the meta-analysis.**

**Table D1: Review of the studies examining liana and tree spectral differences and how they relate with leaf and canopy structure and physiology. L = Liana leaf reflectance, T = Tree leaf reflectance. Results are shown when all examined studies agree on the sign of the difference. When results from several studies diverged, we marked it as inconclusive. The physiological explanation for the differences come directly from Marvin et al. (2016).**

| Spectral region (wavelengths) | Mechanisms controlling reflectance | Relevance to Liana-Tree discrimination | | |
|---|---|---|---|---|
| | | Leaf-level | Canopy level | Physiology |
| Visible (400-700 nm) | Chlorophyll and carotenoid concentrations | L ≥ T | L > T | Lower pigment levels in liana leaves |
| Red Edge (680-700 nm) | Chlorophyll content, water stress, biomass | L ≥ T | L >T | Lower chlorophyll content in liana leaves |
| Near Infrared (700-1400 nm) | Leaf internal structure, nitrogen concentration | T > L | Inconclusive | Physiological differences are inconclusive |
| Shortwave infrared (1500-2500 nm) | Leaf water content, cellulose | Inconclusive | L > T | Leaf water content higher in liana leaves |

### Appendix E: Goodness of fit

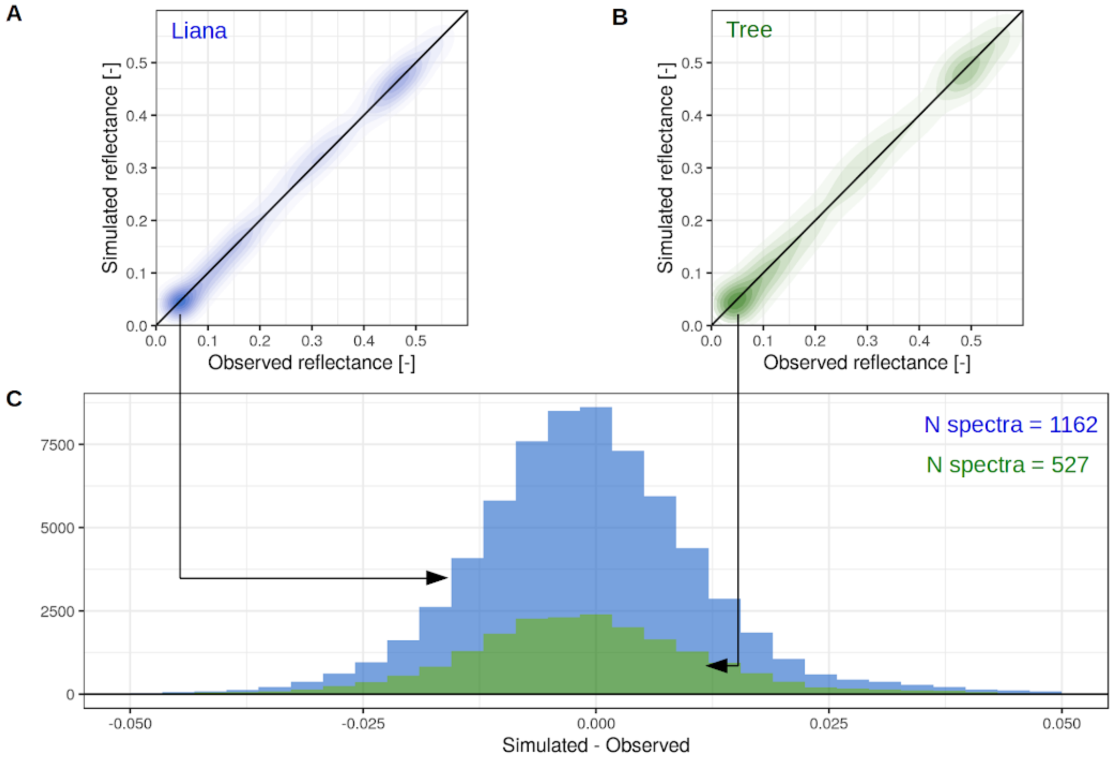


**Figure E1: Liana (blue) and tree (green) goodness of fit for the leaf spectra. In subplots A and B, the point density corresponds to the numbers of observations/simulations (one point per nanometer and species when the raw data were available, one point per nanometer and study/site otherwise). Data points from those subplots were further aggregated in the histograms of subplot C.**

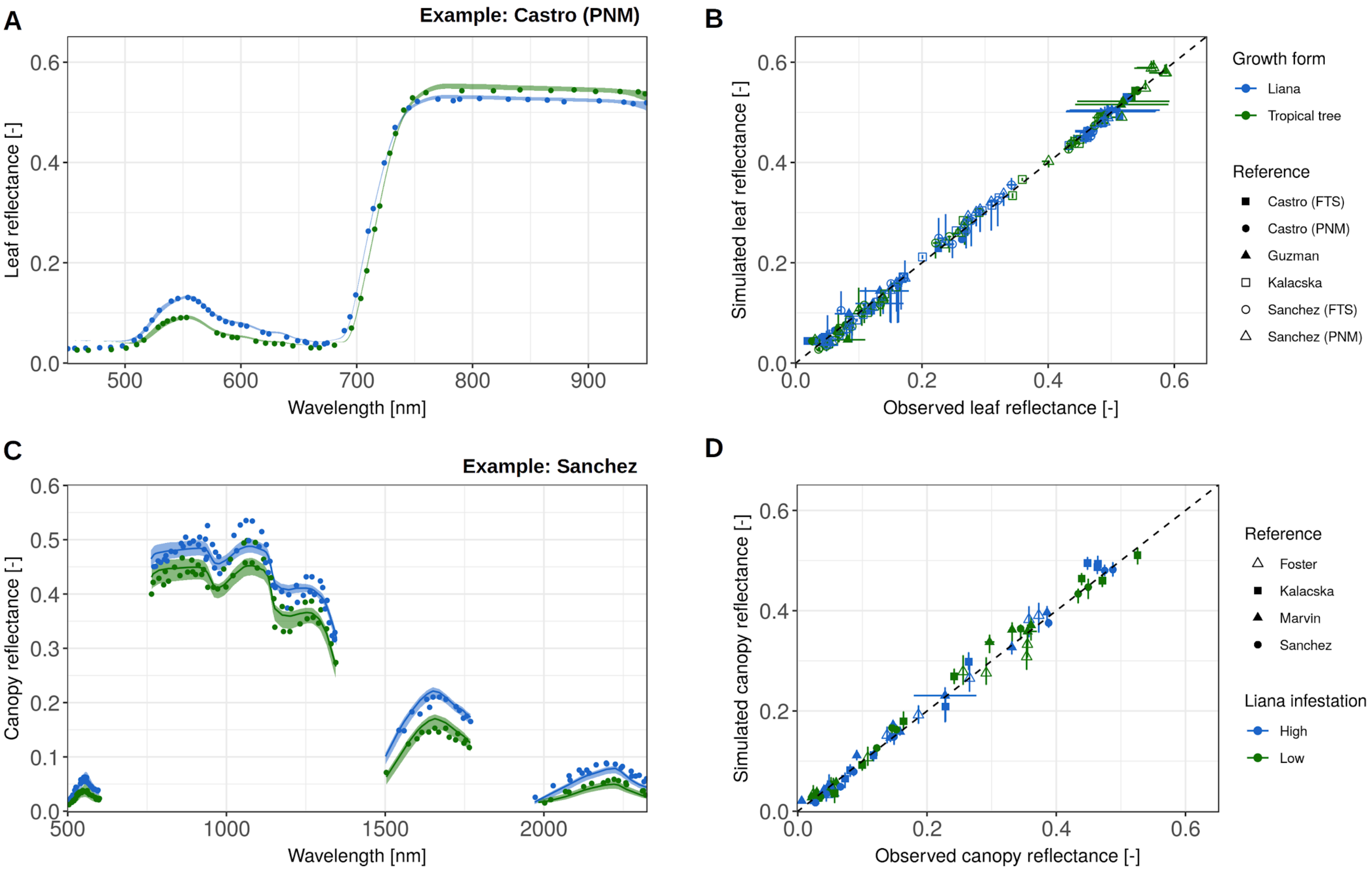


**Figure E2: Leaf and canopy reflectance spectra as observed and simulated by PROSPECT-5 (top) or ED-RTM (bottom), respectively. On the top row, an example of parameter calibration is shown (Castro (PNM), subplot A) alongside with observed *vs* simulated reflectances for all studies together (B). Similarly, on the bottom row, model performance after calibration is illustrated (Sanchez, C) next to the goodness of fit for all studies together (D). In the examples, the points are the measurements while the envelopes encompass the 95% confidence intervals and the solid line the median of the posterior distributions. In B and D, horizontal and vertical error bars represent the observed confidence intervals of data (when communicated) and the prediction intervals, respectively. In these scatterplots, observed and simulated reflectance values were averaged (using a 50 nm window) to make the panel readable.**

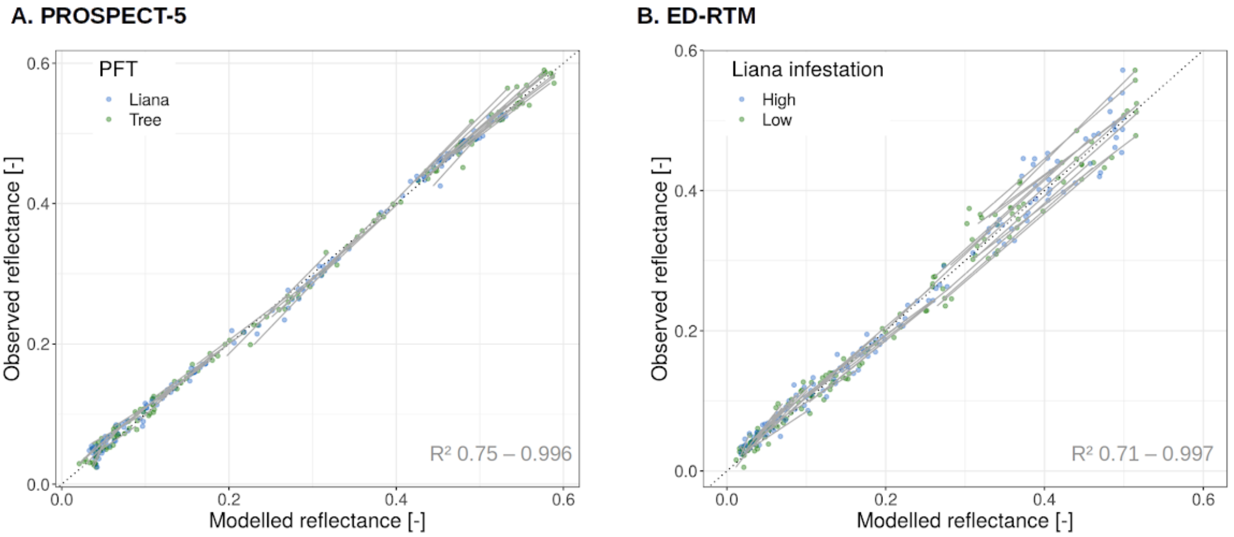


**Figure E3: Observed *vs* modelled leaf (A) and canopy (B) reflectances. In both subplots, values were grouped by wavelength (between 450 and 2500 nm with a 50 nm increment) and the goodness of fit was evaluated with a linear model for each wavelength (grey lines). The range of coefficients of determination for all wavelengths are indicated in the bottom right corner of each subplot.**

**Table E1: Leaf and canopy reflectance spectral error statistics aggregated across the visible (400-700 nm), the near infrared (700-1400 nm) and the shortwave infrared (1500-2500 nm). All metrics, namely the root mean square error (RMSE), the bias (Bias), and the bias-corrected RMSE (SEPC), are unitless (dimension of reflectance). Their mathematical definition is provided in Equation B3.**

| | | | **Visible** | | | **Near infrared (NIR)** | | | **Short wave infrared (SWIR)** | | |
|---|---|---|---|---|---|---|---|---|---|---|---|
| | | | **RMSE** | **BIAS** | **SEPC** | **RMSE** | **BIAS** | **SEPC** | **RMSE** | **BIAS** | **SEPC** |
| **Leaf** | Model vs data | Liana | 0.0064 | 0.0021 | 0.006 | 0.0083 | 0.00067 | 0.0082 | 0.011 | -0.00021 | 0.011 |
| | | Tree | 0.007 | 0.0026 | 0.0065 | 0.006 | 0.001 | 0.0059 | 0.01 | -0.00054 | 0.01 |
| | Data vs data | Liana - Tree | 0.012 | -0.0074 | 0.0098 | 0.02 | 0.019 | 0.0083 | 0.0075 | 0.0001 | 0.0075 |
| | Model vs model | Liana - Tree | 0.012 | -0.0071 | 0.0092 | 0.02 | 0.018 | 0.0079 | 0.0085 | -4.4E-05 | 0.0085 |
| **Canopy** | Model vs data | Liana | 0.016 | -0.0079 | 0.014 | 0.025 | -0.0024 | 0.025 | 0.012 | -0.0012 | 0.012 |
| | | Tree | 0.015 | -0.0075 | 0.013 | 0.026 | -0.00046 | 0.026 | 0.012 | 0.0039 | 0.011 |
| | Data vs data | Liana - Tree | 0.013 | -0.011 | 0.0074 | 0.045 | -0.032 | 0.032 | 0.024 | -0.018 | 0.016 |
| | Model vs model | Liana - Tree | 0.012 | -0.0074 | 0.01 | 0.041 | -0.035 | 0.022 | 0.025 | -0.021 | 0.014 |

### Appendix F: Supplementary results

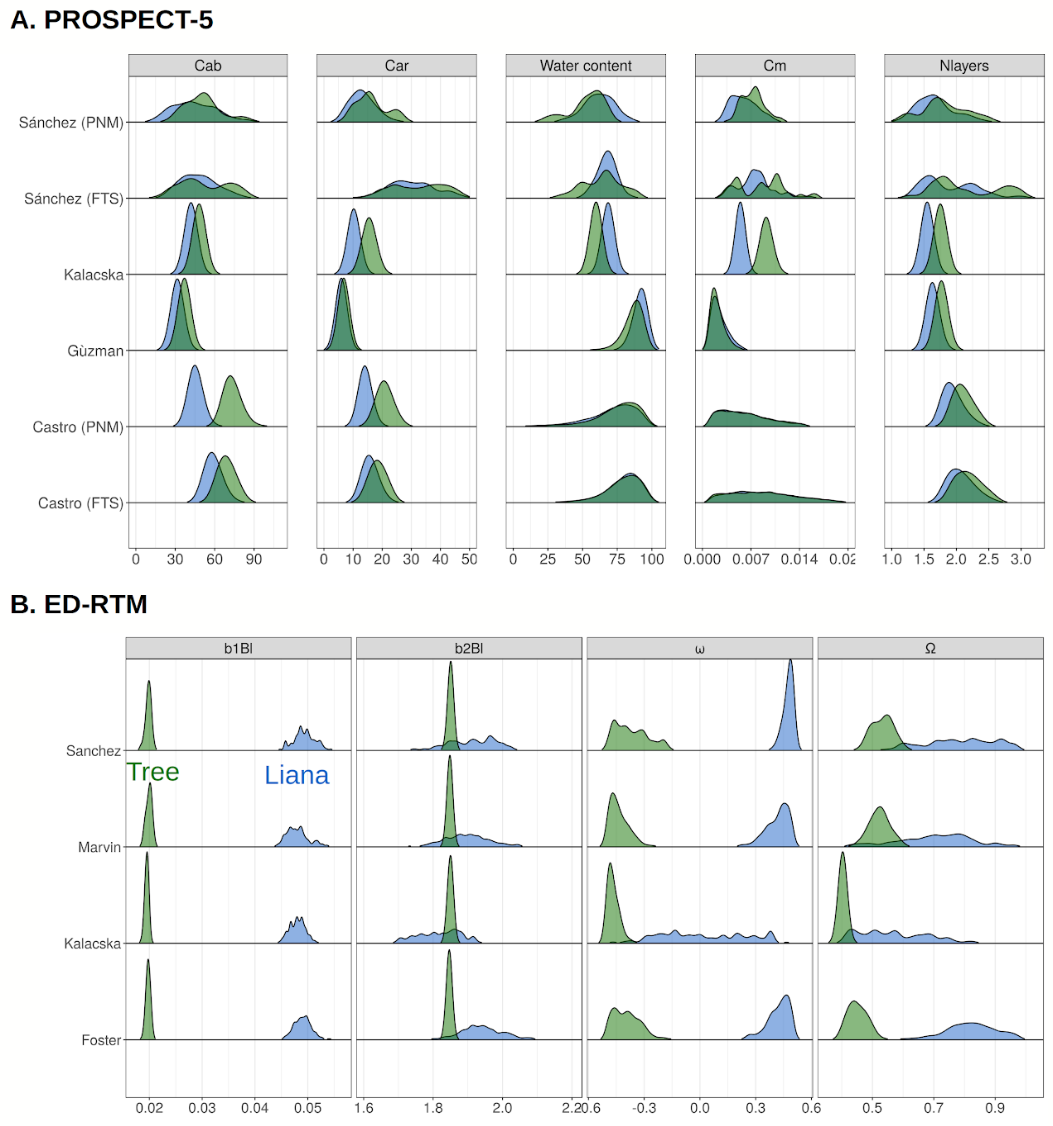


**Figure F1: Posterior distributions of the model parameters after assimilation of the leaf (subplot A) and canopy (subplot B) spectral data for the liana (blue) and the tropical tree (green) PFTs.**

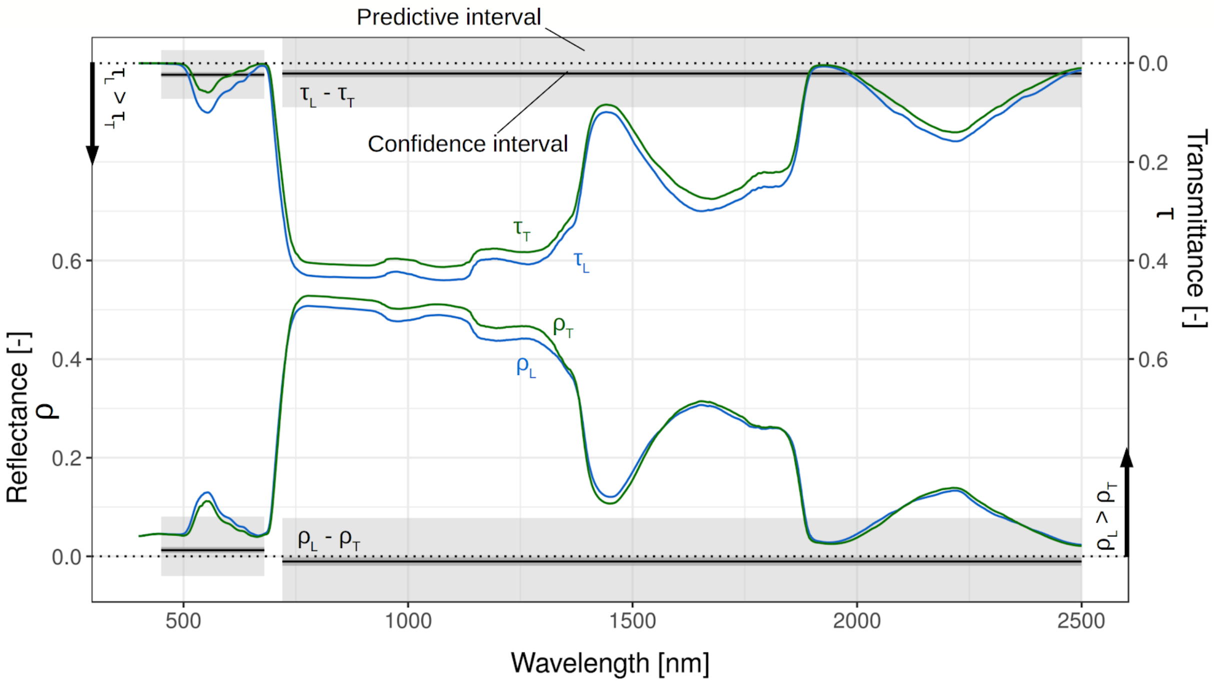


**Figure F2: Liana (blue) and tree (green) leaf spectra, as resulting from the leaf and canopy spectral data calibrations. In subplot A, liana and tree mean reflectance ($\rho_L$ and $\rho_T$, respectively) and transmittance ($\tau_L$ and $\tau_T$, respectively) are plotted alongside with their differences ($\rho_L$ - $\rho_T$ and $\tau_L$ - $\tau_T$) aggregated into broad bands (visible and infrared). The light and dark grey envelopes respectively represent the 95% predictive and confidence intervals of the differences (liana - tree) resulting from 500 liana and tree PROSPECT-5 simulations sampled from the posterior distributions.**

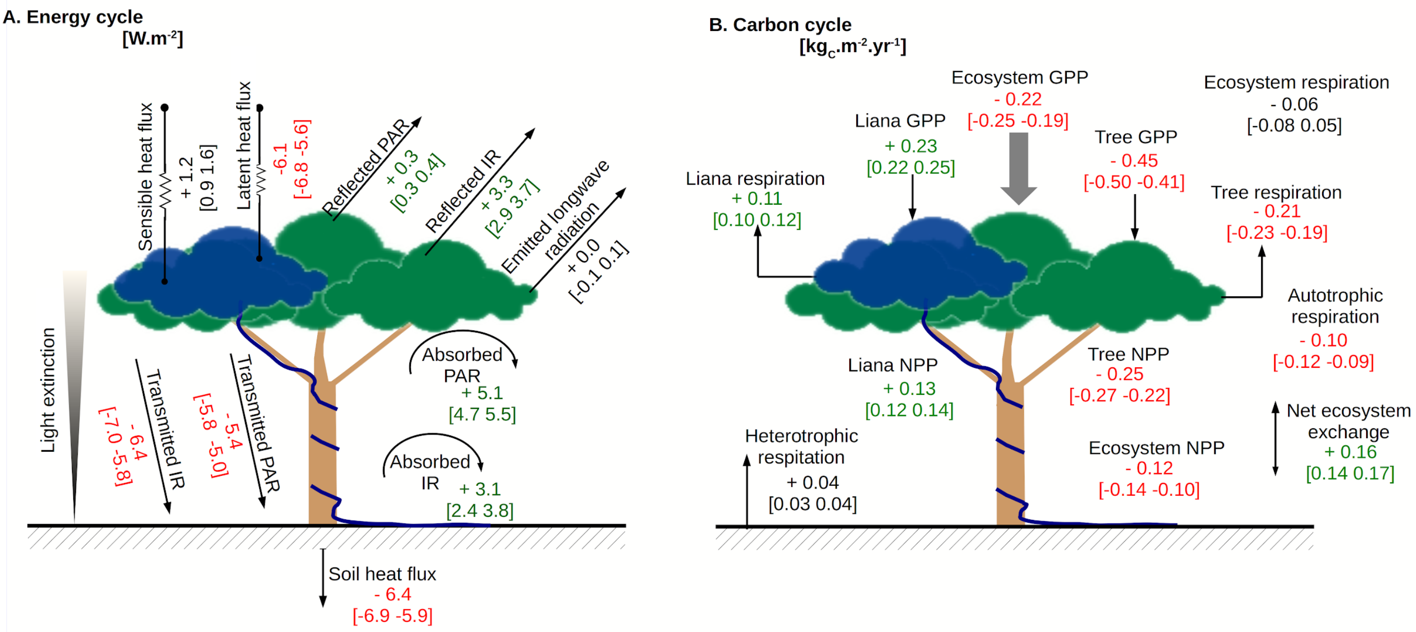


**Figure F3: Mean changes of the energy (A) and Carbon (B) cycle fluxes resulting from the introduction of the liana radiative model parameters together with their confidence intervals. Fluxes are coloured in red (respectively green) when the mean relative changes of the corresponding fluxes are lower than -5% (respectively higher than +5%).**

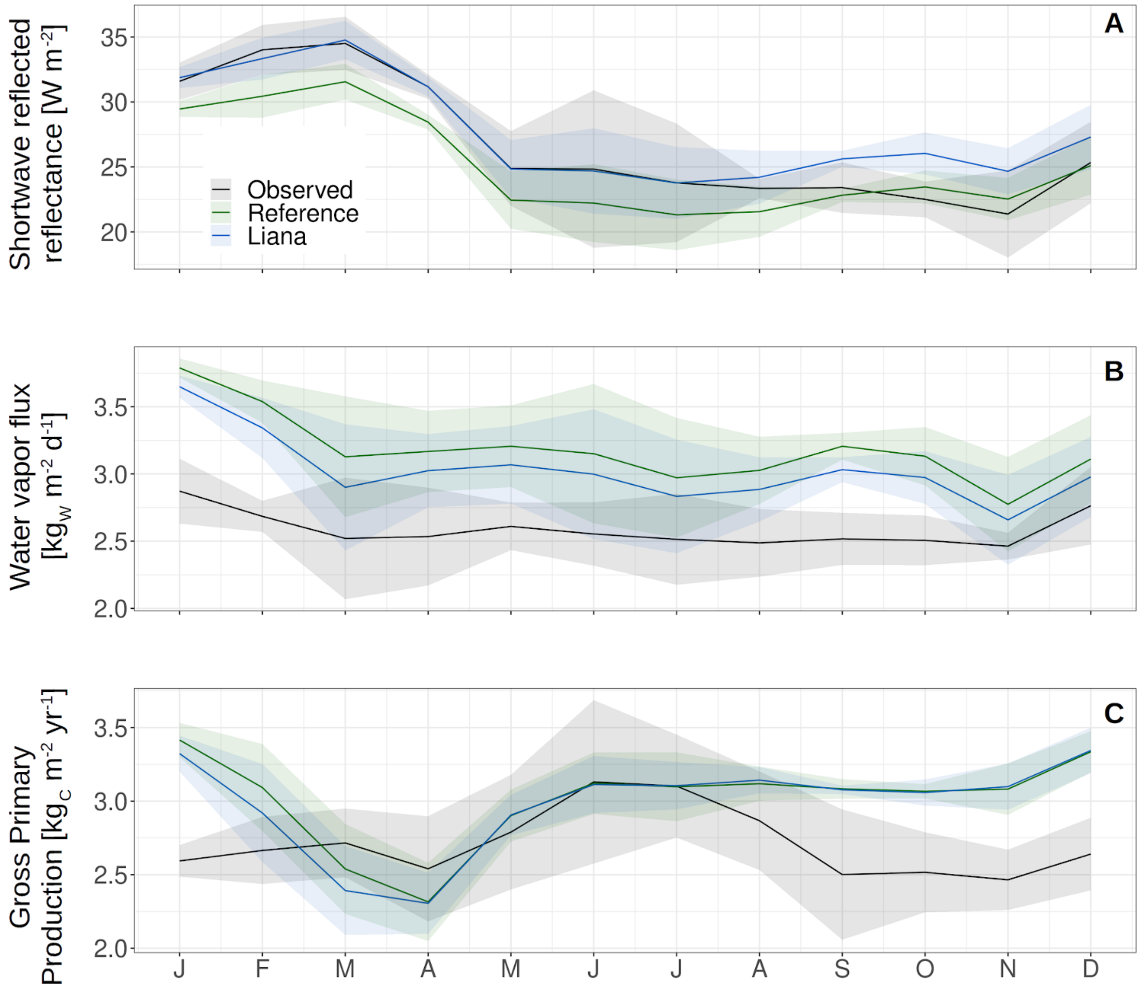


**Figure F4: Simulated (green = "Reference" runs, blue = "Liana" runs) and observed (black) shortwave reflected reflectance (A), latent heat (B), and Gross Primary Production (C). The solid lines represent the mean while the shaded envelopes encompass the interannual variability (mean ± one standard deviation). Including liana optical traits improved the simulated albedo (A), and slightly improved the simulated evapotranspiration (B) and GPP (C) as well.**

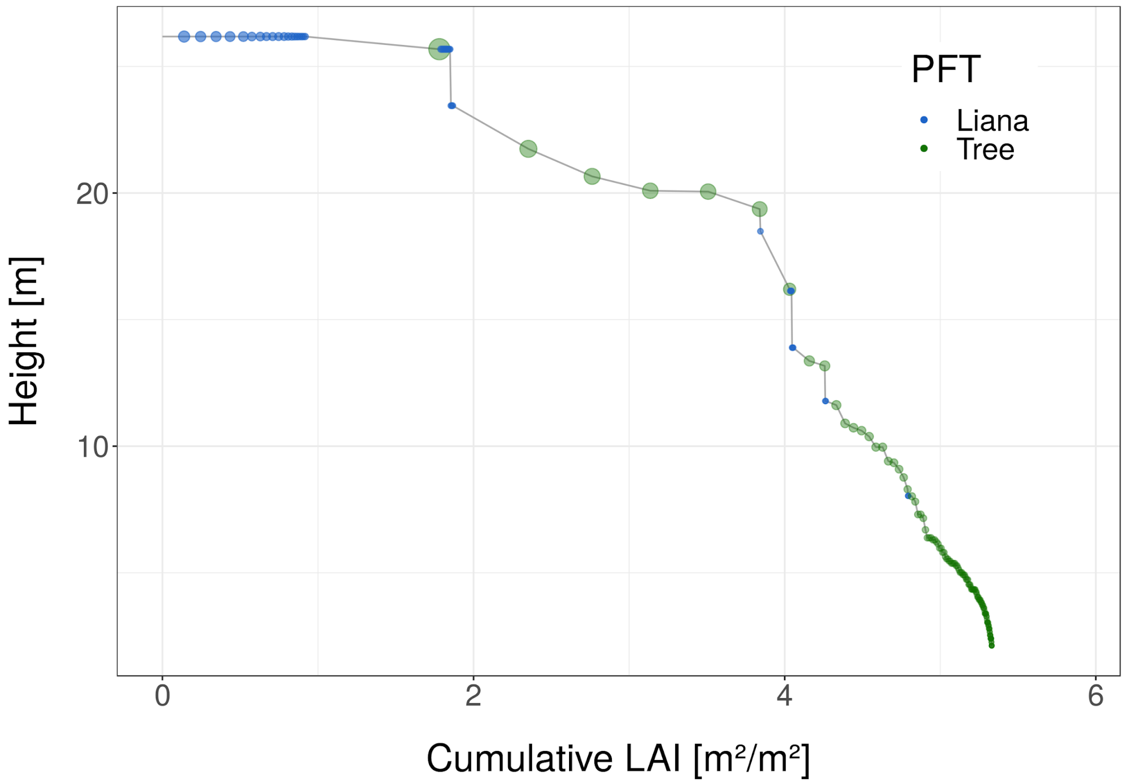


**Figure F5: Example of patch vertical structure on BCI, illustrated as the cumulative LAI (starting from the top canopy) as a function of height. The radius of the data points are proportional to the cohort LAI. Liana vertical clumping is visible at the top of the canopy.**
